# SlS5H silencing reveals specific pathogen-triggered salicylic acid metabolism in tomato

**DOI:** 10.1101/2022.03.03.482652

**Authors:** C Payá, S Minguillón, M Hernández, SM Miguel, L Campos, I Rodrigo, JM Bellés, MP López-Gresa, P Lisón

**Affiliations:** Instituto de Biología Molecular y Celular de Plantas (IBMCP), Consejo Superior de Investigaciones Científicas (CSIC), Universitat Politècnica de València (UPV), Ciudad Politécnica de la Innovación (CPI) 8 E, Ingeniero Fausto Elio s/n, 46011 Valencia, Spain

**Author notes:** **Author for Contact details** Purificación Lisón. **List of author contributions** P.L. conceived the original research plans; I.R., M.P.L.G. and J.M.B. supervised the experiments; M.H. performed the construction of the vectors and the Nicotiana benthamiana experiments, L.C. and P.L. generated the transgenic plants, C.P. and S.M. performed the experiments with Pseudomonas syringae and the senescence studies, C.P. and S.M.M. performed the experiments with CEVd; C.P. performed the experiments with Botrytis cinerea; M.P.L.G. performed the metabolomics analyses; I.R., M.P.L.G. and J.M.B. designed the experiments and analysed the data; P.L. conceived the project and wrote the article with contributions of all the authors; I.R., M.P.L.G. and J.M.B. supervised and completed the writing. P.L. agrees to serve as the author responsible for contact and ensures communication. The author responsible for distribution of materials integral to the findings presented in this article in accordance with the policy described in the Instructions for Authors (www.plantphysiol.org) is: Purificación Lisón.

## Abstract

The phytohormone salicylic acid (SA or 2-hydroxybenzoic acid) plays an important role in plant biotic and abiotic responses. Gentisic acid (GA or 2,5-Dihydroxybenzoic acid, 2,5-DHBA) is the product of the SA 5-hydroxylation which is catalysed by the S5H enzyme, also known as DMR6. GA has been described to accumulate at high levels in compatible plant-pathogen interactions such as tomato plants infected by Citrus Exocortis Viroid (CEVd), and to a much lesser extend upon *Pseudomonas syringae* DC3000 pv. *tomato* (*Pst*) infection. Here we describe the specific effect that tomato SlS5H impairment produces on both plant-pathogen interactions. The induction of *SlS5H* in tomato plants by different pathogens was corroborated by qRT-PCR and correlated with previously described 2,5-DHBA accumulations. Transient *SlS5H* over-expression assays in *Nicotiana benthamiana* confirmed that SA is a substrate for SlS5H *in vivo. RNAi_SlS5H* tomato transgenic plants were generated and characterized upon CEVd and *Pst* infections. Transgenic tomato plants displayed an activation of defences and therefore a loss of susceptibility against both pathogens, and alternative SA homeostasis seems to occur for each specific interaction. Metabolomic assays revealed that whilst the glycosylated form of SA was the most discriminant metabolite found in CEVd infected *RNAi_SlS5H* transgenic plants, trans-feruloyldopamine, feruloylquinic acid, feruloylgalactarate and 2-hydroxyglutarate were the most accumulated compounds in the *Pst*-infected transgenic tomato leaves. Transgenic lines also displayed hyper susceptibility to *Botrytis cinerea*, as well as a smaller size and early senescence. Collectively, our results reveal a novel mechanism by which tomato plants specifically set SA homeostasis upon different pathogen attacks.

**One sentence summary:** The impairment of SA hydroxylation in tomato plants uncovers specific SA homeostasis upon CEVd or *Pseudomonas syringae* infections.

## INTRODUCTION

Salicylic acid (SA or 2-hydroxy benzoic acid) is a phenolic compound present in many plants, and involved in different physiological and biochemical processes, being the activation of inducible defence programs its best characterized function. SA was first described to act in tobacco as an inducer of plant disease resistance to tobacco mosaic virus (White, 1979). Subsequently, evidence suggesting SA is a signal molecule comes from the landmark studies in tobacco mosaic virus-resistant tobacco and cucumber upon infection with necrotizing pathogens (Métraux et al., 1990; Malamy et al., 1992). The essential role of SA in plant defence was definitively demonstrated by using transgenic tobacco plants unable to accumulate SA, which resulted to be incapable to establish the well-known systemic acquired resistance (SAR), an induced defence that confers long-lasting protection against a broad spectrum of pathogens (Gaffney et al., 1993; Delaney et al., 1994). To date, many other studies have been published to point out SA as the best known defence-related hormone (Klessig et al., 2018; Ding and Ding, 2020).

This phenolic compound is biosynthesized in plants from phenylalanine through the route of the phenylpropanoids (PAL pathway) or from isochorismate (IC pathway). Loss-of-function of some genes from both pathways results in an increased plant susceptibility to pathogens, indicating that both the IC and PAL pathways contribute to SA accumulation and function in response to biotic stresses. Nevertheless, the main source of SA when the plant faces a pathogenic infection and SAR is established mostly depends on the IC pathway in *Arabidopsis thaliana* (Wildermuth et al., 2001; Zhang and Li, 2019), being the pathway downstream of IC completely deciphered (Rekhter et al., 2019; Torrens-Spence et al., 2019). In this sense, upon stress situations, ICS1 and ICS2 isochorismate synthase activities isomerise chorismate into IC. In plants, IC is conjugated to the amino acid L-glutamate by an isochorismoyl-9-glutamate (IC-9-Glu) that can spontaneously break down into SA. Besides, EPS1, an IC-9-Glu pyruvoyl-glutamate lyase, can enhance this process more effectively (Zeier, 2021).

Due to its cytotoxic effects, plants maintain SA homeostasis by fine-tuning the balance between the biosynthesis and catabolism of this phytohormone. In this way, SA can be chemically modified into different bio-active derivatives, through glycosylation, methylation, sulfonation, amino acid conjugation, and hydroxylation (Ding and Ding, 2020). Most of the SA present in the plant is glycosylated into SA 2-*O*-β-D-glucoside (SAG) (Enyedi et al., 1992; Malamy et al., 1992) and, to a lesser extent, into salicylate glucose ester (SGE) (Edwards, 1994), being both stored in the vacuole. These conjugates constitute a reserve of inactive SA that can be released slowly in its active form when the plant needs it by the action of glucosyl hydrolases (Dean et al., 2005; Canet et al., 2010). In addition to this, SA can be conjugated with amino acids such as aspartic acid into SA-Asp (Zhang et al., 2007), converted into SA-2-sulfonate by sulfotransferases (Baek et al., 2010), or methylated to form methyl salicylate (MeSA) by salicylic acid carboxyl methyltransferase (SAMT), this latter modification increasing SA membrane permeability and facilitating their mobilization. In SAR, MeSA acts as a phloem-based mobile signal that, after its hydrolysis to SA, triggers resistance (Shulaev et al., 1997; Park et al., 2007).

Regarding SA hydroxylation, a salicylate 3-hydroxylase (AtS3H) was described in Arabidopsis. This hydroxylase is induced by SA and converts this compound into both 2,5-dihydroxybenzoic acid (2,5-DHBA) and 2,3-dihydroxybenzoic acid (2,3-DHBA) *in vitro*, and only into 2,3-DHBA *in vivo*. Studies with *s3h* mutants and the gain-of-function lines revealed that S3H regulates Arabidopsis leaf longevity by mediating SA catabolism (Zhang et al., 2013). DMR6 (Downy Mildew Resistant6) has been proven essential in plant immunity of Arabidopsis (van Damme et al., 2008) and has been later described as a salicylic acid 5-hydroxylase (AtS5H) that catalyses the formation of 2,5-DHBA displaying higher catalytic efficiency than S3H. The Arabidopsis *s5h* mutants and *s5hs3h* double mutants over accumulate SA and display phenotypes such as a smaller size, early senescence, and enhanced resistance to *Pseudomonas syringae* pv. *tomato* DC3000 (Zhang et al., 2017). More recently, the tomato DMR6 orthologs *SlDMR6-1* and *SlDMR6-2* have been identified, also displaying SA 5-hydroxylase activity. Tomato *SlDMR6-1* mutants, obtained by CRISPR/Cas9 system, exhibited broad spectrum disease resistance, correlating this resistance with increased SA levels and transcriptional activation of immune response upon *Xanthomonas gardneri* infection (Thomazella et al., 2021). Therefore, preventing SA hydroxylation confers resistance to pathogens both in Arabidopsis and tomato, standing 2,3-DHBA and 2,5-DHBA for deactivated forms of SA. The main function of these hydroxylated forms is to prevent SA from over-accumulating, thus constituting a mechanism by which plants fine-tune SA homeostasis (Bartsch et al., 2010; Zhang et al., 2013; Zhang et al., 2017; Thomazella et al., 2021).

The 2,5-DHBA or gentisic acid (GA) has been described in animal tissues (Lutwak-Mann, 1943), in microorganisms (Walker and Evans, 1952), and plants (Ibrahim and Towers, 1959). Similar to SA, GA accumulates as glycoconjugates in plants, primarily as GA 5-*O*-β-D-glucosides, GA 5-*O*-β-D-xylosides, or GA 2-*O*-β-D-xylosides (Fayos et al., 2006; Dean and Delaney, 2008; Tárraga et al., 2010; Li et al., 2014). It is also known that exogenous application of GA to tomato (*Solanum lycopersicum*), cucumber (*Cucumis sativus*), and *Gynura aurantiaca* induces the expression of a distinct subset of PR defensive genes compared with the genes induced by SA, as well as mechanisms of gene silencing, and plant resistance (Bellés et al., 1999; Bellés et al., 2006; Campos et al., 2014a).

The accumulation of SA and GA varies up to 100-fold in different species (Zhang et al., 2017). In tomato, that range of difference also occurs among different types of infections, being the accumulation of GA much higher than the SA itself in all of them (Bellés et al., 1999; López-Gresa et al., 2016; López-Gresa et al., 2017; López-Gresa et al., 2019). These differences raise the question whether plants differently regulate homeostasis of SA upon diverse infections. Here we delve into the defensive role of a tomato salicylic acid 5-hydroxylase that provides new insights into the SA metabolism of tomato plants upon different infections.

## RESULTS

### Identification of the tomato pathogen-induced ortholog of *Arabidopsis thaliana S5H*

To identify the enzyme responsible for the conversion of SA into GA in tomato, a Blastp analysis was performed in Sol Genomics databases (https://solgenomics.net/) by using the *AtS5H/DMR6* sequence (At5g24530). A phylogenetic tree was built including the closest tomato sequences (Supplemental Figure S1). The Solyc03g080190 was selected as a candidate for SH5 role in tomato, since it resulted to be one of the closest in the phylogenetic tree, and it presented the highest identity percentage (67.35%), in contrast to the 62.43% of identity displayed by the Solyc06g073080 sequence, also near in the phylogenetic tree. The Blastp analysis of the Solyc03g080190 in The Arabidopsis Information Resource (https://www.arabidopsis.org/) confirmed the selected candidate, being *AtS5H/DMR6* the closest gene to Solyc03g080190 *(SlS5H)* in *Arabidopsis thaliana*. Finally, this sequence coincided with At5g24530 tomato ortholog proposed by *EnsemblePlants* (http://plants.ensembl.org/index.html) and with *SlDMR6-1*, which has been recently proposed as the *DMR6* ortholog in tomato and which displayed salicylic acid 5-hydroxylase activity (Thomazella et al., 2021).

According to available transcriptome data, *SlDMR6-1* expression has been described to be induced in response to several pathogens (Thomazella et al., 2021). To confirm the pathogen triggered *SlS5H* induction, and to extend the study of expression to other pathogens which provoke the accumulation of SA and GA, tomato plants were subjected to infection either with Citrus Exocortis Viroid (CEVd), Tomato Spotted Wilt Virus (*TSWV*), Tomato Mosaic Virus (*ToMV*) or a virulent and an avirulent strain of *Pseudomonas syringae* pv. *tomato* DC3000 (*Pst*) (see Materials and Methods). Leaf samples were collected at the indicated time points and *SlS5H* expression levels were analysed by qRT-PCR in those tomato-pathogen interactions (Figure 1A to 1D). A clear induction of *SlS5H* was observed upon all the pathogen infections in the analysed samples, reaching levels up to 5 times higher in CEVd-infected or *Pst-*infected plants than those observed in the non-infected plants, and around 2 times higher in the case of TSWV or ToMV infections. It is worthy to note that this induction patterns correlated with symptomatology, causing infection with CEVd the most severe disease symptoms and being ToMV infection symptomless (López-Gresa et al., 2012). Regarding the bacterial infection, *SlS5H* expression levels were higher in the tomato plants inoculated with the virulent bacteria (*Pst* DC3000 ΔAvrPto) as compared to the avirulent infection at 48 hours post inoculation (hpi), therefore confirming this tendency.

**Figure 1.**
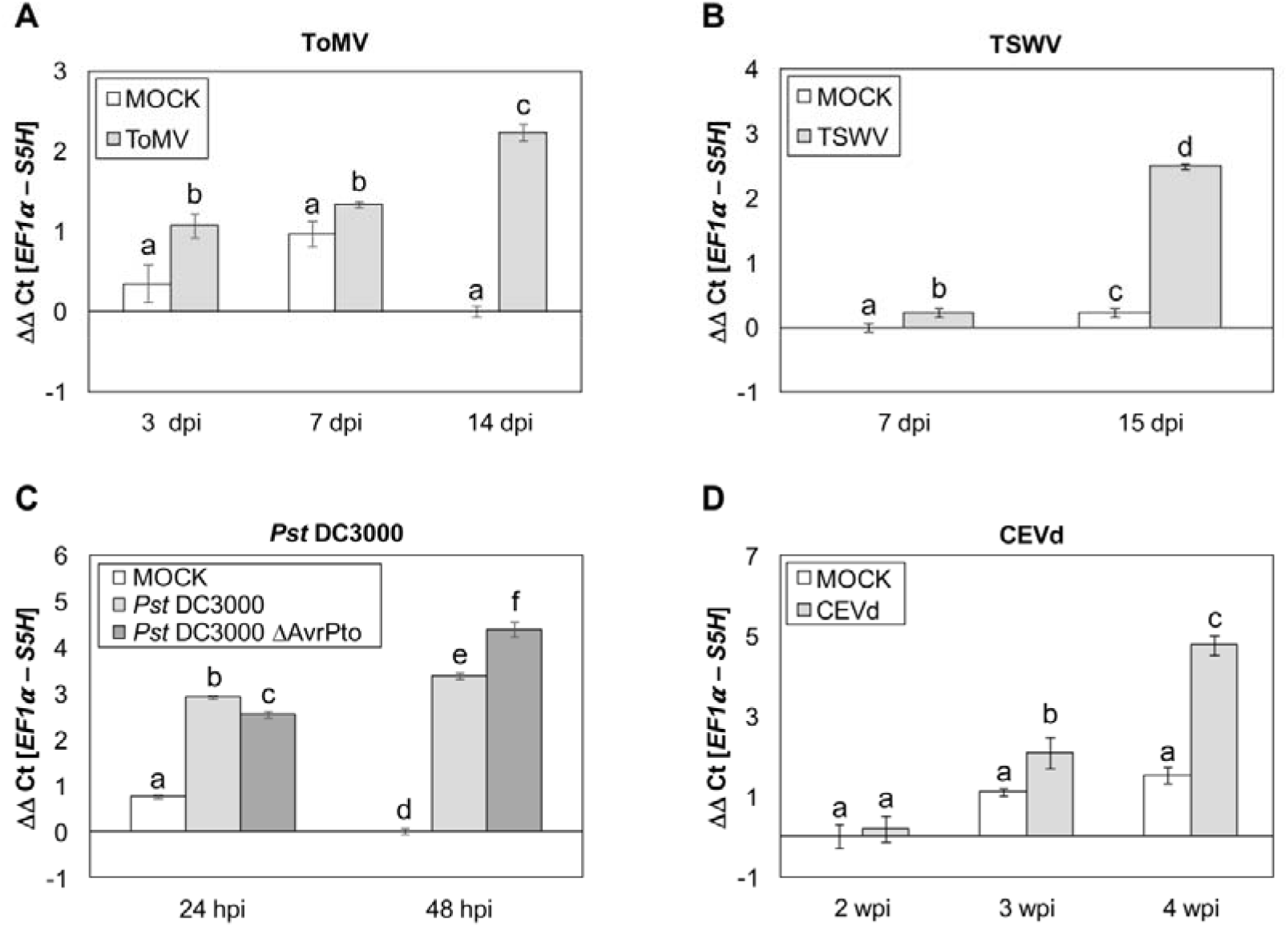
*SlS5H* expression patterns induced by different pathogens. Expression of *SlS5H* gene after inoculation with Tomato Mosaic Virus (ToMV, **A**); Tomato Spotted Wilt Virus (TSWV, **B**); *Pseudomonas syringae* pv. *tomato* DC3000 (*Pst*DC3000 **C**) and Citrus Exocortis Viroid (CEVd, **D**) at different times post-inoculation. MOCK represents the mock-inoculated plants.

To study the induction of *SlS5H* by its own substrate, SA treatments were performed, and samples were collected at different time points (see Materials and Methods). As Figure S2A shows, a statistical induction of *SlS5H* was detected by qRT-PCR at 6 hours post treatment (hpi) when compared with mock inoculated plants, presenting the maximum induction at 1 hpt. *PR1* activation was used as a positive control for the treatments, presenting a significant induction at 24 hpt (Figure S2B).

All these data confirm that *SlS5H* is involved in the plant response to pathogens, extending its putative role to different tomato-pathogen interactions.

### Overexpression of *Sl5H* in *Nicotiana benthamiana* decreases SA levels *in vivo*

To confirm S5H biochemical activity *in vivo, Nicotiana benthamiana* plants were agroinoculated with a construction carrying the *SlS5H* cDNA containing a His tag under the *35S CaMV* promoter. These plants (*pGWB8-SlS5H*) and the corresponding control plants inoculated with the empty plasmid (*pGWB8*) were then embedded with SA (see Materials and Methods). The accumulation of the recombinant protein was confirmed by western blot analysis in *pGWB8-SlS5H* plants, and levels of free and total SA were measured (Figure S3). As expected, levels of free and total SA were almost 3 times lower in *pGWB8-SlS5H* plants, being these differences statistically significant, and thus confirming SA is a substrate for SlS5H *in vivo*. However, no differences in neither the GA nor in 2,3-DHBA accumulations were detected between pGWB8 and pGWB8-SlS5H *Nicotiana benthamiana* plants (Figure S3D).

### Silencing *SlS5H* increases resistance to CEVd in tomato

To gain further insights into the role of SlS5H *in vivo*, silenced transgenic Moneymaker tomato plants were generated by following an RNAi strategy (see Materials and Methods). The generated tomato lines *RNAi_SlS5H* were characterized, and several independent transgenic lines were confirmed. Homozygous lines *RNAi_SlS5H 14* and *RNAi_SlS5H 16* both carrying one copy of the transgene, were selected for further studies.

To extend the role of *SlS5H* in plant defence, the tomato-CEVd interaction was selected, since GA -the proposed product of S5H activity-had been described to accumulate at very high levels in CEVd-infected tomato plants (Bellés et al., 1999; López-Gresa et al., 2016; López-Gresa et al., 2017; López-Gresa et al., 2019). Therefore, wild type (WT) and *RNAi_SlS5H* transgenic plants were inoculated with CEVd and checked for the development of symptoms. The characteristic symptomatology of CEVd-infected tomato plants consists of epinasty, stunting, leaf rugosity, midvein necrosis and chlorosis (Prol et al., 2020). As Figure 2A shows, differences in the percentage of plants displaying symptoms were observed in both *RNAi_SlS5H* transgenic lines with respect to the parental tomato plants. Particularly, transgenic line *RNAi_SlS5H 16* displayed only 35% of plants showing symptoms at 1.9 weeks post inoculation (wpi), while almost 75% of non-transgenic plants exhibited them. Moreover, at 2.3 wpi all the WT plants displayed symptoms while around 20% of the *RNAi_SlS5H* remained symptomless. These results indicate a delay in symptom appearance in *RNAi_SlS5H* tomato plants.

**Figure 2.**
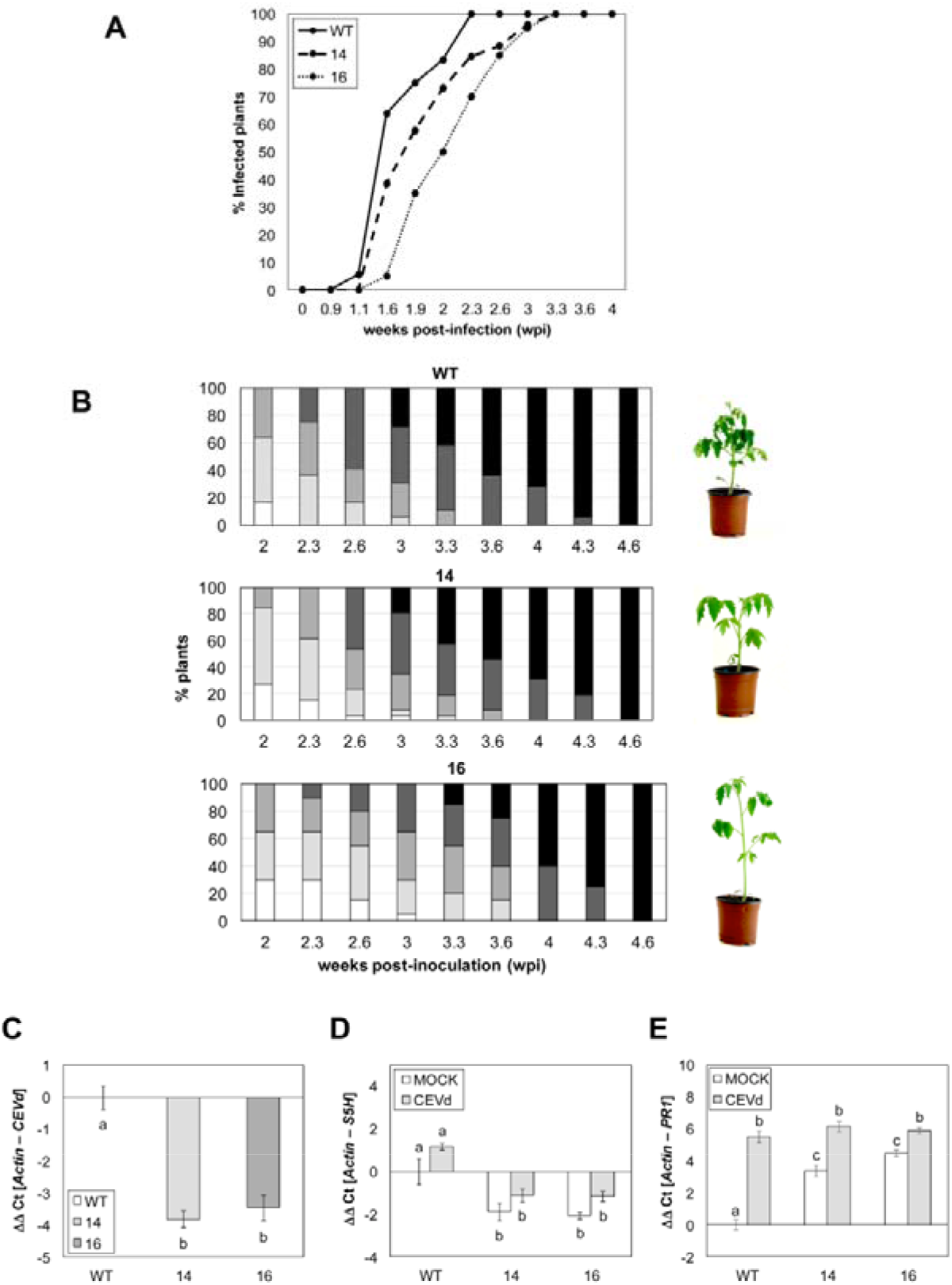
Symptomatology and transcriptomic analysis in wild type (WT) and *RNAi_SlS5H* (lines 14 and 16) tomato plants infected with CEVd. **(A)** Disease development in WT and *RNAi_SlS5H 14* and *RNAi_SlS5H 16* plants infected with CEVd. Evolution of the percentage of tomato plants showing symptoms at the indicated time points (wpi). **(B)** Representative images and disease severity of CEVd infected WT and *RNAi_SlS5H* plants. Symptomatology was established using the following scale: no symptoms (white), mild epinasty (light grey), severe epinasty and stunting (grey), leaf rugosity (dark grey), midvein necrosis and chlorosis (black). Data correspond to one representative experiment. **(C)** CEVd accumulation in WT and transgenic lines in CEVd-infected plants three weeks post inoculation (wpi). *S5H* **(D)** and *PR1* **(E)** tomato genes were analyzed 2 weeks after inoculation with CEVd. Expression levels are normalized to the tomato actin gene. Data correspond to the mean ± standard error of technical and biological replicates of a representative experiment (n=3). Different letters indicate statistically significant differences (*p* < 0.05).

To confirm the differences in the disease development observed, a scale of the disease severity was established, scoring symptoms from mild (mild epinasty) to very severe (midvein necrosis and chlorosis), at different time points (see Materials and Methods). As Figure 2B shows, differences were observed between WT and *RNAi_SlS5H* transgenic tomato plants from 2.3 to 3.6 wpi. Moreover, *RNAi_SlS5H 16* transgenic line did not display very severe symptoms at 3 wpi, whilst 30% parental plants exhibited severe symptoms at the same time point. Therefore, the observed differences in symptom severity confirmed the loss of susceptibility of *RNAi_SlS5H* tomato plants to CEVd infection. To confirm the enhanced resistance, the presence of pathogen was measured at 3 weeks post-inoculation (wpi), detecting a statistically significant decrease in the CEVd accumulation in both *RNAi_SlS5H* transgenic lines (Figure 2C).

Our results appear to indicate *S5H* silencing reduces tomato susceptibility to CEVd, confirming the role of *SlS5H* in the plant defence response.

### Silencing *SlS5H* causes an activation of plant defence upon CEVd infection

To confirm that *SlS5H* expression levels in the RNAi transgenic plants infected with CEVd were downregulated throughout infection, qRT-PCR from WT, *RNAi_SlS5H 14* and *RNAi_SlS5H 16* plants, were performed at 2 weeks after the inoculation with CEVd (Figure 2D). Effectively, levels of *SlS5H* expression were significantly lower in both transgenic lines before (mock) and upon viroid infection (CEVd) than the corresponding wild type (WT) tomato plants. To find out if *RNAi_SlS5H* transgenic lines exhibited an activation of the defensive response against CEVd, the expression of the pathogenesis related protein 1 (*PR1*; accession X71592), which has been described as a classical marker of plant defence rapidly induced in CEVd-infected tomato plants (Granell et al., 1987; Tornero et al., 1993), was also studied by qRT-PCR at 2 wpi. As expected, *PR1* was induced by CEVd in WT plants, but also in both *RNAi_SlS5H* transgenic lines (Figure 2E). Interestingly, levels of expression of *PR1* were already higher in mock-infected *SlS5H*-silenced tomato plants. These results appear to indicate that *SlS5H* silencing provokes an activation of the SA-mediated plant defence.

To better characterise the plant response of the *RNAi_SlS5H* transgenic lines upon CEVd infection, the expression of several defence genes was also studied at 3 wpi both in mock and infected plants. Regarding the gene silencing mechanisms which are activated by CEVd in wild type plants, a significant reduction in *DCL1* and *DCL2* induction was observed in CEVd-infected *RNAi_SlS5H* lines when compared to the corresponding wild type plants, thus indicating a correlation between the amount of CEVd and the induction of these two dicers (Figure S4A and S4B). However, no significant differences were observed in the *RDR1* induction pattern in the transgenic plants, which displayed a significant induction of this gene upon viroid infection (Figure S4C). As far as jasmonic acid (JA) response is concerned, a statistically significant reduction of the JA-induced proteinase inhibitor *TCI21* (Lison et al., 2006), was observed in both mock-inoculated *RNAi_SlS5H* lines when compared with wild type (Figure S4D), indicating that the JA final response is repressed in these tomato transgenic plants.

Therefore, we have observed that *RNAi_SlS5H* transgenic lines display constitutive *TCI21* repression as well as *PR1* induction in response to CEVd infection, thus suggesting a possible increase of SA levels in these transgenic lines.

### Levels of SA and GA are altered in *RNAi_SlS5H* transgenic tomato plants upon CEVd infection

Free and total levels of SA and its hydroxyl product GA were quantified in the wild-type and *RNAi_SlS5H* transgenic lines upon viroid infection at 3 wpi (Figure 3). As expected, free and total SA and GA levels were higher in all the CEVd-infected plants.

**Figure 3.**
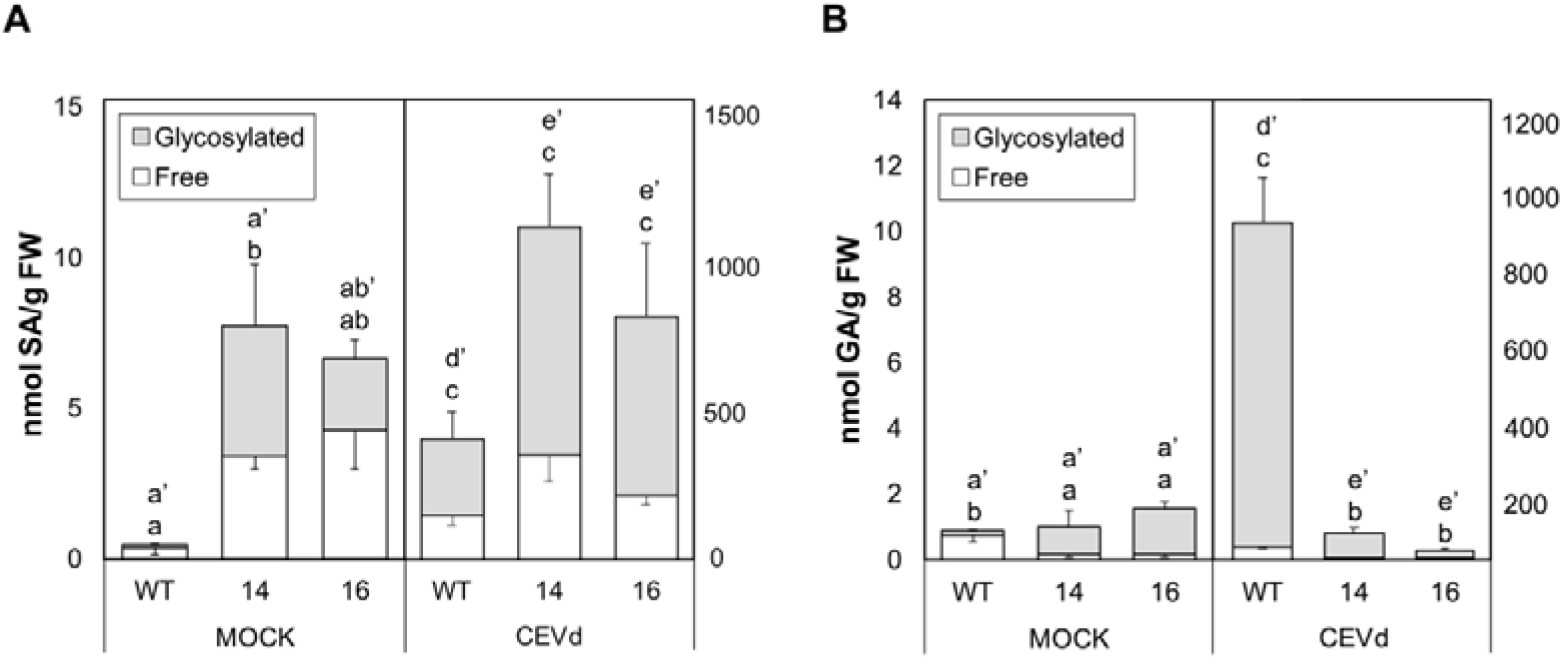
Accumulation of free and glycosylated SA (A) and GA (B) in wild type (WT) and transgenic *RNAi_SlS5H* tomato plants infected with CEVd. Phenolic compounds were extracted from infected leaves 3 weeks after inoculation; MOCK represents the mock-inoculated plants. The extracts were analyzed by fluorescence HPLC. Bars represent the mean ± standard deviation of a representative experiment (n=4). Significant differences in free and glycosylated SA and GA accumulation between lines and infected or mock-inoculated plants are represented by different letters since *p-*value < 0.05

In CEVd-inoculated plants, levels of total SA in both *RNAi_SlS5H* transgenic infected lines resulted to be significantly higher when compared with those observed in control infected plants, reaching 1000 nmol/g fresh weight whilst levels in control infected plants barely reached 400 nmol/g fresh weight (Figure 3A). SA levels in non-pathogenic conditions (mock) were also higher in both transgenic lines, being free SA levels in transgenic *RNAi_SlS5H* 14 line statistically significant.

Once detected the over-accumulation of SA, we decided to study the levels of GA, which is the product of the S5H activity. As Figure 3B shows, a drastic reduction of total GA levels was observed upon viroid-infected in both *RNAi_SlS5H* transgenic lines as compared to the levels detected in the infected WT tomato plants (10-fold). Interestingly, this significant reduction was also observed in free GA corresponding to mock conditions.

The higher levels of SA and the lower accumulation of GA found in the CEVd-infected *RNAi_SlS5H* transgenic lines confirm the decrease of the salicylate 5-hydroxylase activity *in vivo* and explain the observed enhanced resistance and activation of the SA-mediated plant defence in these transgenic plants.

### Infection with *Pseudomonas syringae* pv. *tomato* DC3000 produces specific phenolic accumulation pattern in *RNAi_SlS5H* transgenic tomato plants

Bacterial infection of tomato plants produces lower accumulation of SA (López-Gresa et al., 2017). To study the role of *SlS5H* in this tomato-pathogen interaction, WT and *RNAi_SlS5H* transgenic tomato plants were infected with the virulent strain of *Pst*, and bacterial counting from infected leaves was carried out at 24 hours after infection (hpi). As shown in Figure 4A, a significant 10-fold decrease in the number of colony forming units was observed in infected tissues of the different transgenic lines with respect to their genetic background, confirming that transgenic plants *RNAi_SlS5H* also exhibited resistance against *Pst*.

**Figure 4.**
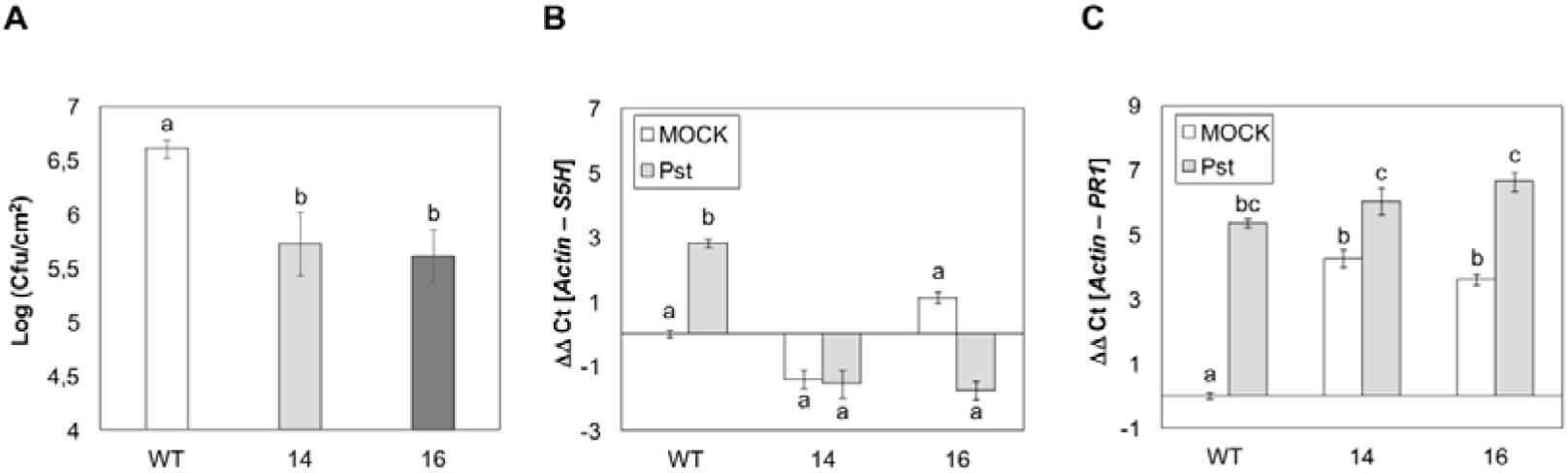
Growth of *Pseudomonas syringae* pv. *tomato* DC3000, and *S5H* and *PR1* gene expression in leaves of wild type (WT) and 14 and 16 lines of *RNAi_SlS5H* transgenic plants. Tomato plants were inoculated with *Pst* DC3000 by immersion (OD_600_ = 0.1). Leaf samples for bacterial growth quantification **(A)**, and *S5H* **(B)** and *PR1* **(C)** gene expression analysis were taken 24 hours after bacterial infection. MOCK represents the non-inoculated plants. The qRT-PCR values were normalized with the level of expression of the actin gene. The data are presented as means ± standard deviation of a representative experiment (n=3). Statistically significant differences (*p*-value < 0.05) between genotypes and infected or mock-treated plants are represented by different letters.

*SlS5H* silencing in tomato plants upon bacterial infection was also confirmed by qRT-PCR, being differences in *SlS5H* expression in bacterial infected *RNAi_SlS5H* transgenic plants statistically significant compared to the expression levels observed in the infected WT (Figure 4B). Similar to what was observed upon viroid infection, *PR1* expression was already higher in mock-inoculated SlS5H-silenced tomato plants and was significant induced by bacteria in both WT and *RNAi_SlS5H* transgenic lines (Figure 4C).

In a similar manner to that performed with CEVd-infected plants, SA and GA levels were measured in both WT and *RNAi_SlS5H* transgenic plants upon bacterial infection (Figure 5). In *Pst*-tomato interaction, the levels of free and total SA were induced by the biotrophic pathogen in all the analysed plants. Strikingly unlike what was observed upon CEVd infection, the bacteria produced a significant reduction of free and total SA levels in both *RNAi_SlS5H* transgenic lines when compared with levels observed in *Pst*-infected WT plants (Figure 5A). Regarding GA, only WT tomato plants showed a statistical accumulation of total GA levels after bacterial infection, thus indicating that the product of *SlS5H* is also reduced in *RNAi_SlS5H* transgenic plants upon bacterial infection (Figure 5B).

**Figure 5.**
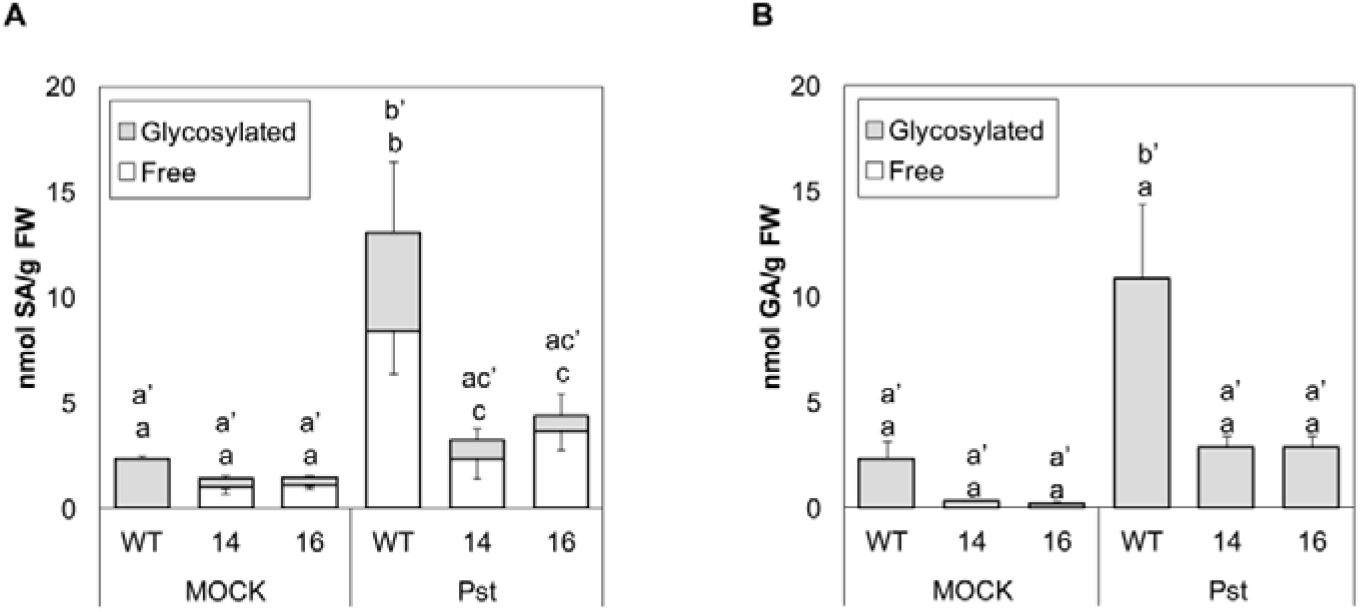
Accumulation of free and glycosylated SA (A) and GA (B) in wild type (WT) and transgenic *RNAi_SlS5H* (lines 14 and 16) tomato plants infected with *Pseudomonas syringae* pv. *tomato* DC3000. Phenolic compounds were extracted from infected leaves 24 hours post-inoculation; MOCK represents the mock-inoculated plants. The extracts were analyzed by fluorescence HPLC. Bars represent mean ± standard deviation of a representative experiment (n=4). Significant differences on free and glycosylated SA and GA accumulation between genotypes and infected or mock-inoculated plants are represented by different letters since *p-* value < 0.05

The reduction of SA levels observed in *RNAi_SlS5H* transgenic lines upon bacterial infection appears to indicate that SA is specifically catabolized upon pathogen infection in these transgenic plants.

### A metabolomic analysis of the *RNAi_SlS5H* transgenic tomato plants upon viroid and bacterial infection reveals differences in SA metabolism upon pathogen attack

To better understand the SA metabolism in *RNAi_SlS5H* transgenic lines upon each infection, a metabolomic study based on ultra-performance liquid chromatography-mass spectrometry (UPLC-MS) was performed. For viroid infection, 8-day-old tomato plants were used, and samples were collected 3 weeks after CEVd inoculation (wpi), while bacterial infection was carried out on 5-week-old tomato plants and the harvesting time was 24 h after *Pst* infiltration (hpi). Then, hydro alcoholic extracts from control and infected *RNAi_SlS5H* tomato leaves were analysed by UPLC-MS, and multivariate data analysis was employed to deal with the large number of mass data. Specifically, a principal component analysis (PCA) was first applied to identify metabolic changes after viroid and bacterial infection of tomato plants (Figure 6A). An extensive separation between both tomato interactions was observed by PC1 due to the different experimental conditions: temperature and plant age. Reaching the PC3, the metabolic changes in tomato leaves produced by both pathogens was explained, being greater those caused by viroid inoculation. To exclude the differences due to this biological variability, two different PCA were therefore applied separately on each tomato-interaction.

**Figure 6.**
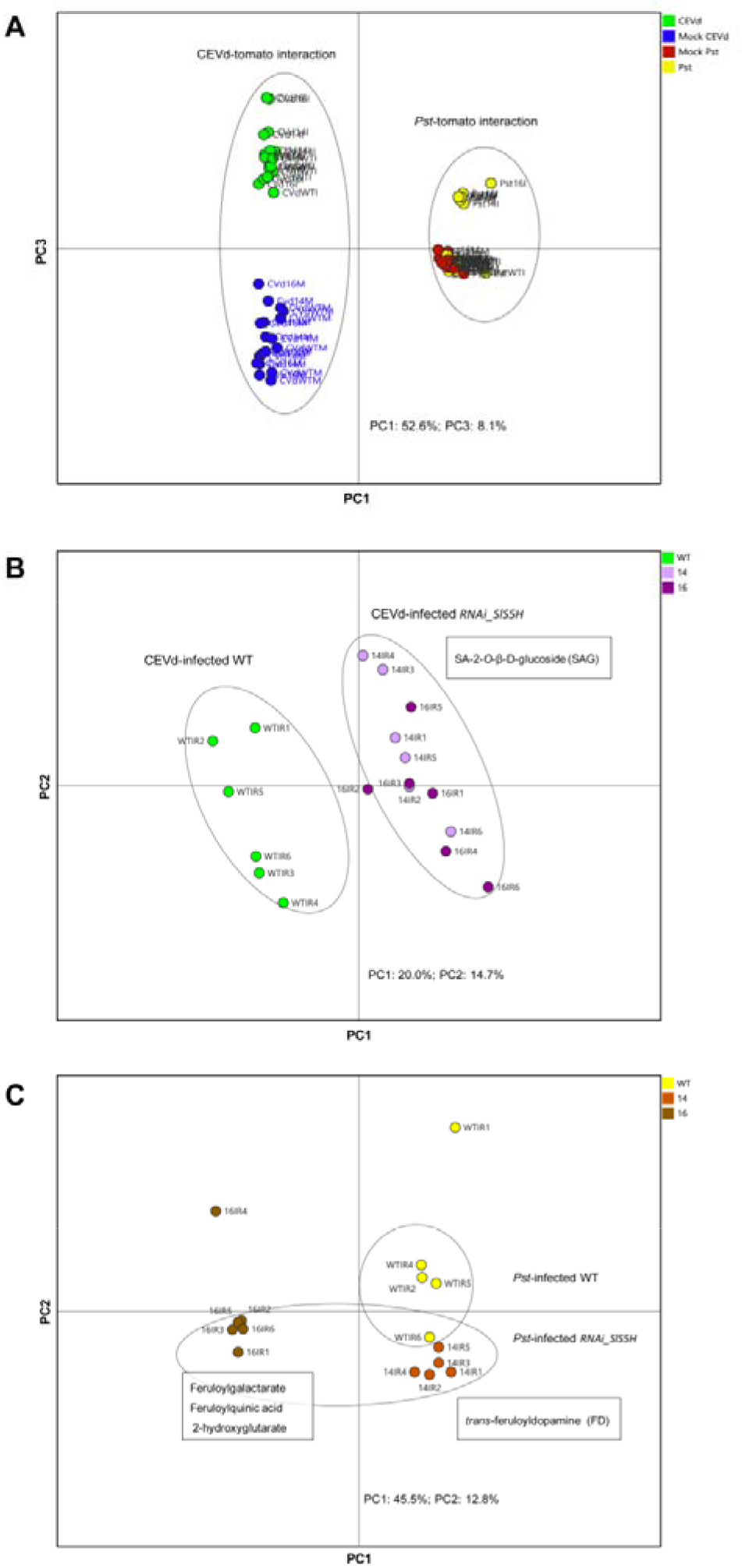
PCA Score plots based on whole range of on the whole array of the mass spectra within a *m/z* range from 100 to 1500 using unit variance (UV) scaling method of methanolic extracts from tomato leaves. **(A)** Green: Infected by CEVd (3 wpi); Blue: Mock for CEVd infection; Yellow: Infected by *Pst* (24 hpi); Red: Mock for bacterial infection. **(B)** Green: Wild type (WT); light purple: CEVd infected *RNAi_SlS5H* 14 at 3 wpi; dark purple: CEVd infected *RNAi_SlS5H* 16 at 3 wpi. **(C)** Yellow: Wild type (WT); orange: *Pst* infected *RNAi_SlS5H* 14 at 24 hpi; brown: *Pst* infected *RNAi_SlS5H* 16 at 24 hpi.

In particular, the first two components of the PCA score plot of viroid-tomato interaction (Figure S5A) divided the observations by the infection (mock *vs*. infected plants; PC1) and genotype (WT *vs*. transgenic plants; PC2). In order to elucidate the SA metabolism in tomato plants against CEVd, a PCA of both infected wild type (WT) and transgenic *RNAi_SlS5H* plants was performed (Figure 6B). For the identification of the metabolites accumulated in the infected *SlS5H* silenced lines (14 and 16), the positive PC1 loading plot was analysed. Interestingly, the glycosylated form of SA (SAG) was the most accumulated compound in the transgenic plants (fold change transgenic lines *vs*. WT: 3.0; *p*-value 0,009). These results are in accordance with the total SA accumulation measured by HPLC-fluorescence in CEVd infected *RNAi_SlS5H* plants (Figure 3A).

In the case of bacteria-tomato interaction, the PC3 of PCA (Figure S5B) explained the different metabolic content of transgenic tomato plants from WT, while PC1 clearly discriminated the metabolome of the infected *RNAi_SlS5H* line 16. In a similar manner to CEVd interaction, to investigate the role of SA in the bacterial infected tomato plants, the PCA of infected tomato plants was required (Figure 6C). By analysing the negative PC1 and PC2 loading plot, the metabolites accumulated in both transgenic lines were identified. In contrast to CEVd infection, SAG accumulation was not induced by *Pst* in both transgenic lines according to the results obtained using fluorescence-based detection (Figure 5A). In tomato-bacteria interaction, some of the compounds over-accumulated in the transgenic infected leaves were identified as feruloyldopamine (fold change between transgenic lines and WT: 5.3; *p*-value 0.003), feruloylquinic acid (fold change: 5.1; *p*-value 0.006), feruloylgalactarate (fold change: 3.4; *p*-value 0.01) and 2-hydroxyglutarate (fold change: 1.4; *p*-value 0.04).

### *SlS5H* silencing reveals differences in SA biosynthesis gene expression upon pathogen attack

To study differential expression of genes participating in SA biosynthesis, qRT-PCR were performed for *ICS, PAL, EPS1* and *SAMT* genes in samples corresponding to CEVd or *Pst* infections for both wild type and *RNAi_SlS5H* transgenic plants (Figure 7).

**Figure 7.**
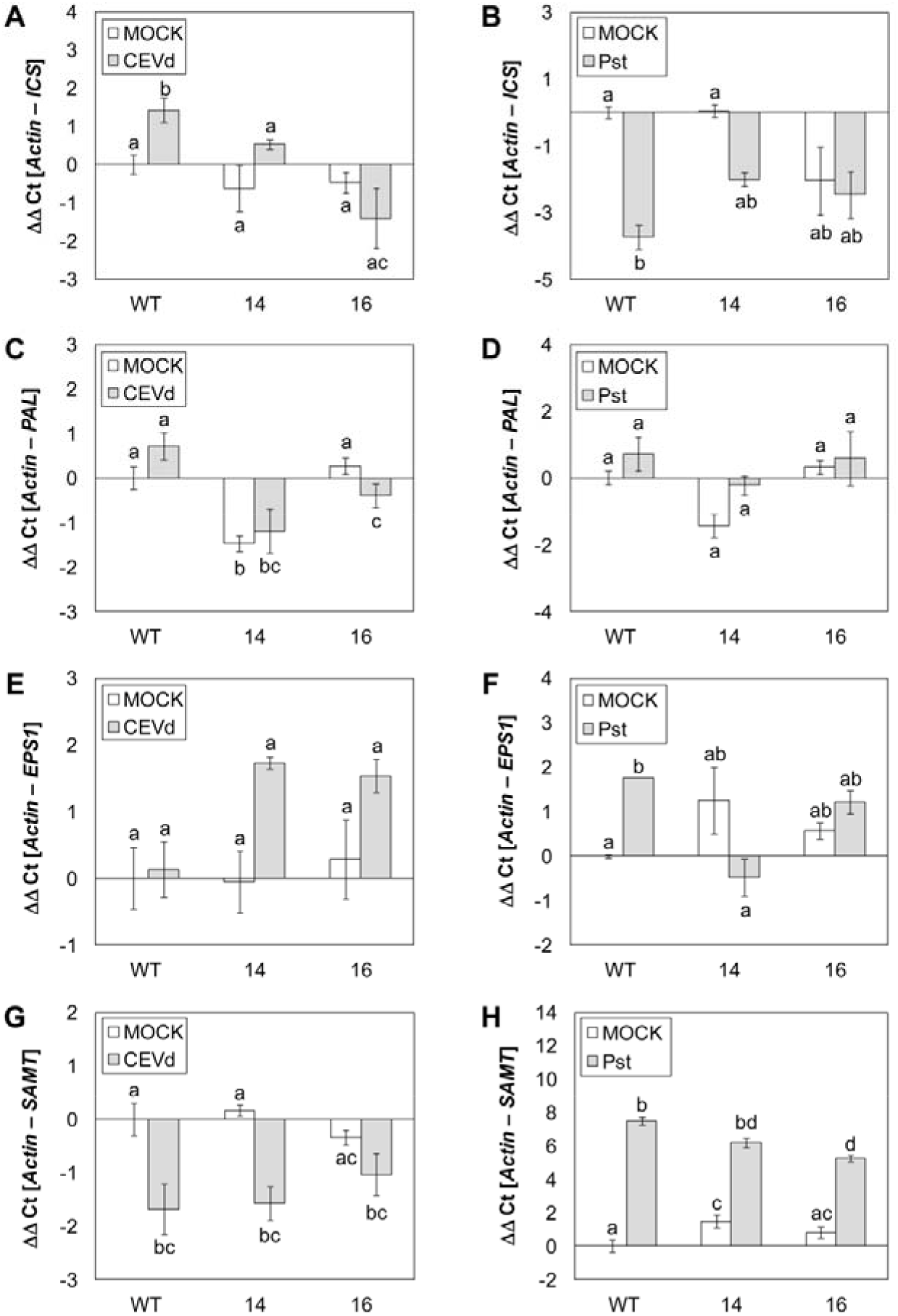
Differences in the pattern of induction of SA metabolism genes upon viroidal (CEVd) and bacterial (Pst) inoculation in wild type (WT) and *RNAi_SlS5H* (lines 14 and 16) transgenic tomato plants by qRT-PCR. *ICS* **(A)**, *PAL* **(C)**, *EPS1* **(E)** and *SAMT* **(G)** tomato gene expression levels 3 weeks post-inoculation with CEVd; *ICS* **(B)**, *PAL* **(D)**, *EPS1* **(F)** and *SAMT* **(H)** gene expression levels 24 hours after inoculation with the bacteria *Pseudomonas syringae* DC3000; MOCK represents the mock-inoculated plants. The qRT-PCR values were normalized with the level of expression of the actin gene. The expression levels correspond to the mean ± the standard error of a representative experiment (n=3). Statistically significant differences (*p-* value < 0.05) between genotypes and infected or mock-inoculated plants are represented by different letters.

As Figure 7A shows, a significant induction in *ICS* was observed in wild type plants upon CEVd infection, being that induction impaired in the transgenic lines, thus suggesting that the SA biosynthesis is down-regulated when the SA hydroxylation is prevented, which may lead to SA over-accumulation. Contrasting with CEVd infection, a clear reduction of *ICS* expression was detected upon bacterial infection, in both WT and transgenic lines. However, this decrease of expression was not statistically significant in the transgenic plants (Figure 7B).

As far as *PAL* pattern of expression is concerned, whilst CEVd infection caused a significant reduction in both transgenic lines compared to infected WT, no significant differences caused by *Pst* infection were observed (Figures 7C and 7D).

The last step in the SA biosynthesis through the isochorismate pathway involves the conversion of isochorismate-9-glutamate into SA, being performed by *EPS1*. Whilst wild type plants appear not to display any induction of *EPS1* by CEVd, bacterial infection clearly provoked the induction of this gene. This pattern was completely opposite in both *RNAi_SlS5H* transgenic lines, since they showed a slight induction upon CEVd infection, being impaired in the expression of *EPS1* upon bacterial infection (Figures 7E and 7F).

Finally, a reduction of *SAMT* expression upon CEVd infection was measured in all the genotypes, with no clear differences observed between WT and transgenic plants. In contrast, *SAMT* induction caused by bacterial infection was reduced in transgenic plants, showing statistically significant differences between the induction observed in infected WT and *RNAi_SlS5H 16* transgenic plants (Figure 7H).

Although the differences observed in both viroid and bacterial infections were not statistically significant for all the analysed genes, a noticeable variation in the pattern of expression of SA biosynthesis genes provoked by these two pathogens was detected, thus suggesting SA metabolism is specific for the tomato-pathogen interaction.

### *SlS5H* silencing causes repression of the JA defence response

Once confirmed that *SlS5H* silencing provokes an activation of the SA-mediated defence, we studied the possible cross-talk with JA response. To that purpose, control and *RNAi_SlS5H* transgenic lines were wounded and the induction of *TCI21* was studied at 24 h after wounding (Figure 8A). As expected, *TCI21* was highly induced by wounding in wild type plants, whilst *RNAi_SlS5H* wounded plants displayed only a slight induction of TCI21, which were comparable to non-wounded plants, thus indicating that JA response is repressed in these transgenic plants. Interestingly, a higher induction of *PR1* was detected in the transgenic plants both in non-wounded and upon wounding when compared with the WT plants (Figure 8B), reinforcing the observed activation of SA-mediated defence in the *RNAi_SlS5H* transgenic plants.

**Figure 8.**
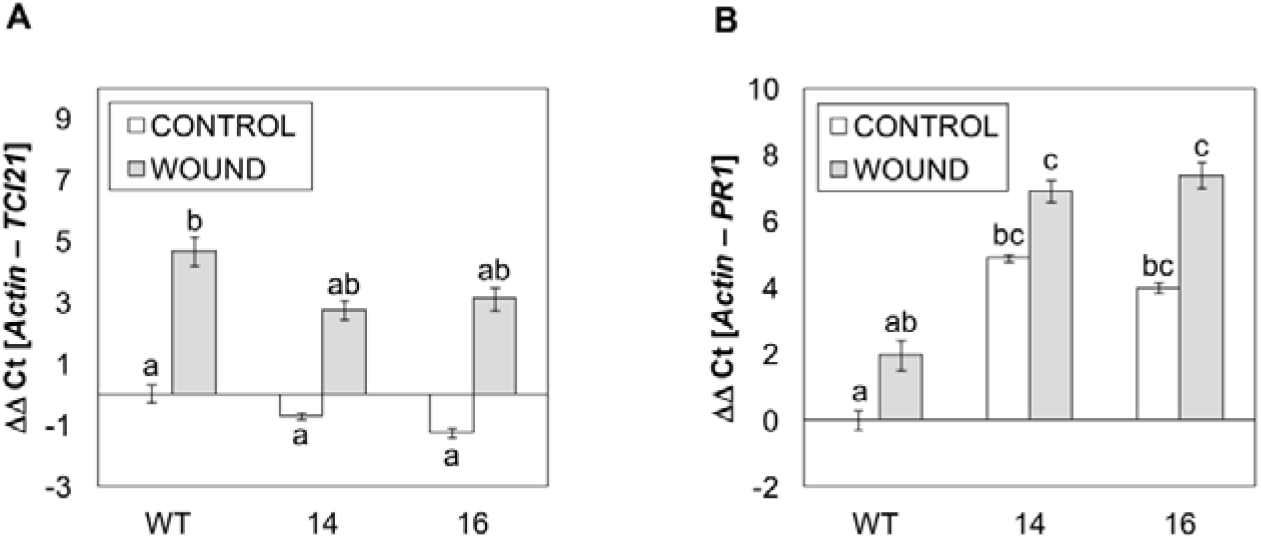
Expression of the tomato *TCI21* (A) and *PR1* (B) genes in wild type plants (WT) and the *RNAi_SlS5H* (lines 14 and 16) transgenic tomato plants in control (CONTROL) and wounded (WOUND) plants. Samples were taken 24 h after wounding; CONTROL represents the non-wounded plants. The qRT-PCR values were normalized with the level of expression of the actin gene. The expression levels correspond to the mean ± the standard error of a representative experiment (n=3). Statistically significant differences (*p-*value < 0.05) between genotypes and infected or mock-inoculated plants are represented by different letters.

Since JA response appeared to be repressed in *RNAi_SlS5H* transgenic lines, the phenotype against the necrotrophic fungal pathogen *Botrytis cinerea* was explored (Figure 9A). Although no statistically significant differences were found in the size of necrotic lesions between the transgenic lines and the corresponding parental plants (Figure 9B), both *RNAi_SlS5H* transgenic lines displayed an increase in the susceptibility against *Botrytis cinerea*, as indicated by the increase in the yellowish area shown by the transgenic plants. To better quantify this effect, chlorophyll content was measured in control and transgenic plants infected with *Botrytis cinerea*. As Figure 9C shows, *RNAi_SlS5H* transgenic lines accumulated significant lower levels of chlorophyll B and total chlorophylls, therefore confirming the observed increased susceptibility of *RNAi_SlS5H* transgenic lines against *Botrytis cinerea*.

**Figure 9.**
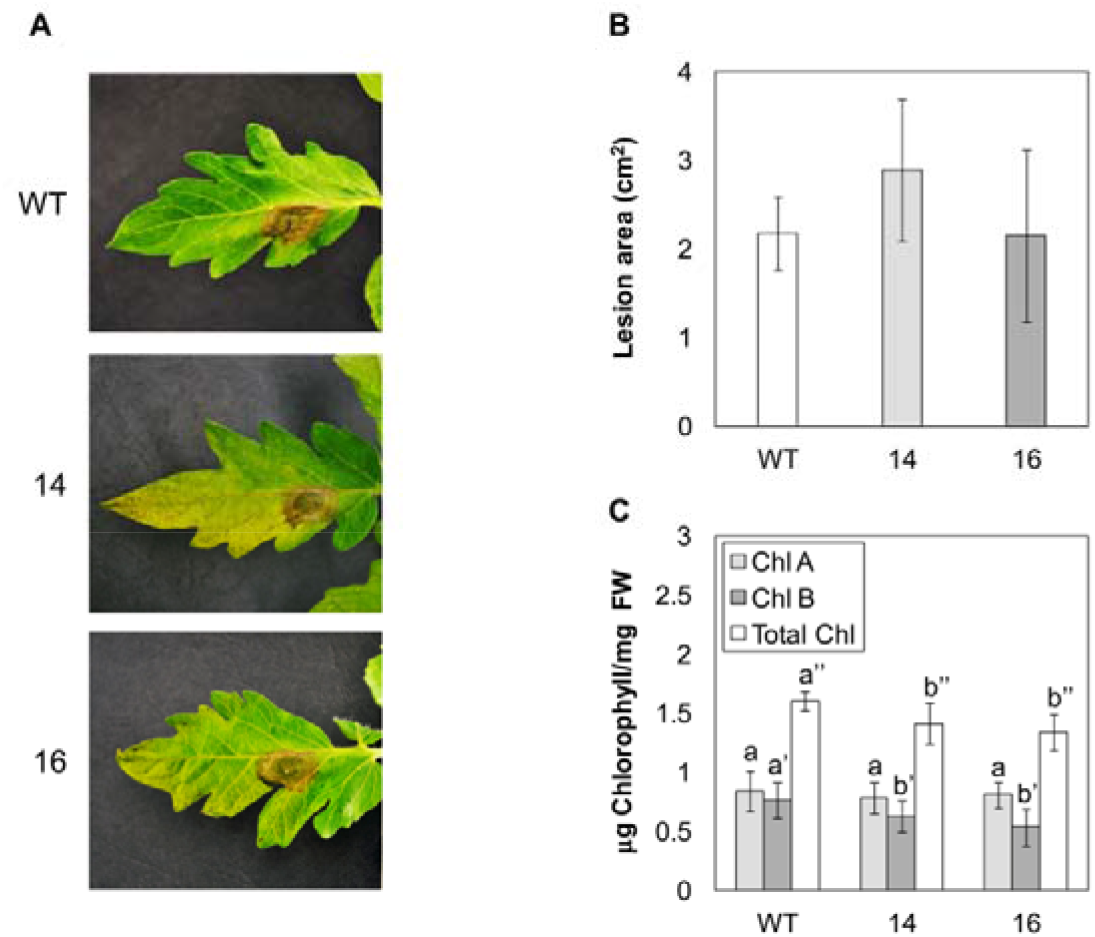
Response of wild type plants (WT) and *RNAi*_S5H transgenic lines 14 and 16 to *Botrytis cinerea* infection. Representative images **(A)**, lesion area **(B)** and chlorophyll content **(C)** of plants infected with *B. cinerea* 5 days after fungal inoculation. Bars represent the mean ± the standard deviation of a representative experiment (n=6). Significant differences between genotypes are represented different by letters since *p-*value < 0.05

### Silencing of *SlS5H* results in early senescence

Previous research on salicylate hydroxylase activity in Arabidopsis reported that the deficiency in this enzyme provokes an advanced senescent response (Zhang et al., 2013; Zhang et al., 2017). However, no noticeable phenotypic differences were described in *SlDMR6-1* tomato mutants (Thomazella et al., 2021).

To study the phenotype of *RNAi_SlS5H* transgenic tomato plants, 5 plants for each genotype were grown for 10 weeks, and the percentages of leaflets displaying different senescence stages were recorded. As Figure 10A shows, *RNAi_SlS5H* transgenic plants displayed and earlier chlorosis, even leading to the leaf collapse. This change in colour associated with senescence is due to the disappearance of chlorophylls which reveals the other underlying photosynthetic pigments such as carotenes and xanthophylls (Junker and Ensminger, 2016). Particularly, in 10-week-old plants we observed that control plants still possess approximately 50% of the green leaves, while hardly any leaf of the transgenic lines remained green, turning the entire observed leaves to different intensities of yellow and brown, eventually leading to leaf fall (Figure 10B).

**Figure 10.**
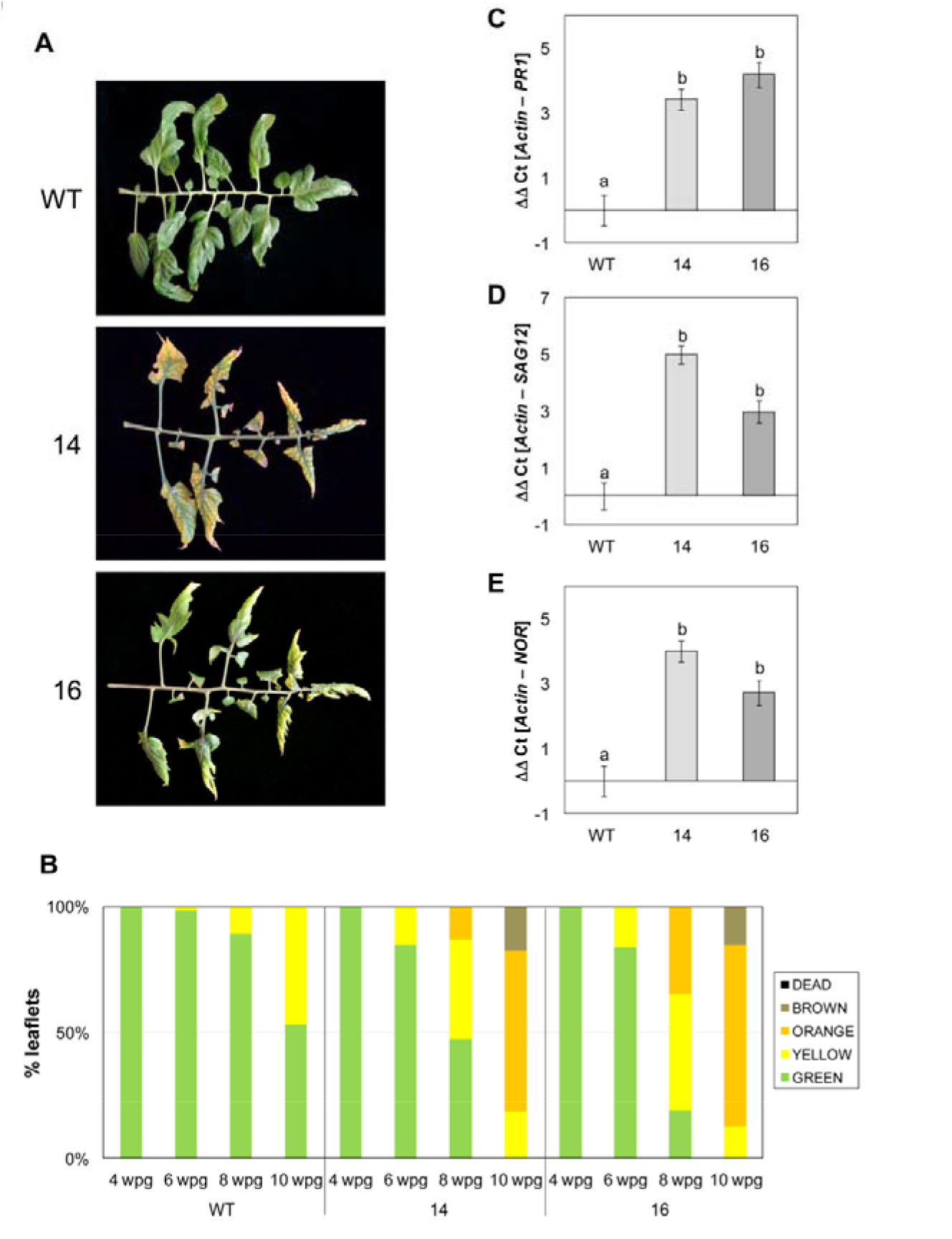
Early senescent phenotype and senescence-related gene expression analysis in WT and *RNAi*_S5H transgenic lines 14 and 16. **(A)** Representative images and **(B)** evolution of senescence in WT and *RNAi_S5H* tomato leaflets. The percentage of leaflets of same appearance with respect to the total of leaflets corresponding to the third, fourth and fifth leaves of the WT and *RNAi_SlS5H* transgenic tomato plants at 4, 6, 8 and 10 weeks post-germination (wpg) is shown. “GREEN” refers to the natural color of the tomato leaves, “YELLOW” when the leaves begin to age and acquire yellowish spots, “ORANGE” is the time when the leaf turns completely yellow, “BROWN” when necrotic lesions appear due to aging and “DEAD” defines those leaves that have detached from the plant or that are completely necrotic. Expression levels of *PR1* **(C)**, *SAG12* **(D)** and *NOR* **(E)** are shown for WT plants and *RNAi_SlS5H* transgenic lines 14 and 16, 4 weeks after germination. The qRT-PCR values were normalized with the level of expression of the actin gene. The expression levels correspond to the mean ± the standard error of a representative experiment (n=3). Statistically significant differences (*p-*value < 0.05) between genotypes are represented by different letters.

To better characterize the phenotype of the *RNAi_SlS5H* transgenic tomato plants, we measured weight and conductivity at 10 weeks after germination, observing a significant weight reduction in the transgenic plants (Figure S6A). Moreover, electrolyte leakage, which is a hallmark of cell death, was also significantly increased in both transgenic lines silencing *SlS5H* (Figure S6B). Finally, the chlorophyll content was also measured again observing significant lower levels of chlorophyll B and total chlorophylls in *RNAi_SlS5H* transgenic tomato leaves (Figure S6C).

To reinforce the data obtained by phenotypic visualization, a gene expression analysis for *PR1* as well as the senescence markers *SAG12* (AT5G45890 tomato ortholog; Solyc02g076910) and *NOR* (Ma et al., 2019) was carried out by qRT-PCR, at 4 weeks after germination. As shown in Figure 10, a significant expression of *PR1* was observed in *RNAi_SlS5H* transgenic lines with respect to control plants, which could correlate with the higher SA levels previously observed in mock-inoculated plants (Figure 5A). In parallel with *PR1* expression levels, both senescence markers *SAG12* and *NOR* were differentially upregulated in the *RNAi_SlS5H* transgenic leaves, confirming the phenotype of early senescence proposed for these plants.

Levels of free and total SA and GA were measured at different stages of development (Figure 11). To that purpose, samples were collected at the indicated time points and free and total levels of both phenolics were measured by HPLC in both control and *RNAi_SlS5H* transgenic tomato plants. In general, levels of SA were higher in the transgenic plants impaired in its hydroxylation, being differences with control plants in total SA statistically significant at 10 weeks after germination. However, levels of GA resulted to be lower at any time point, being total GA levels statistically significant from 6 weeks after germination, which confirms that the reduction of GA in these transgenic plants is consistent and reproducible in all the analysed samples.

**Figure 11.**
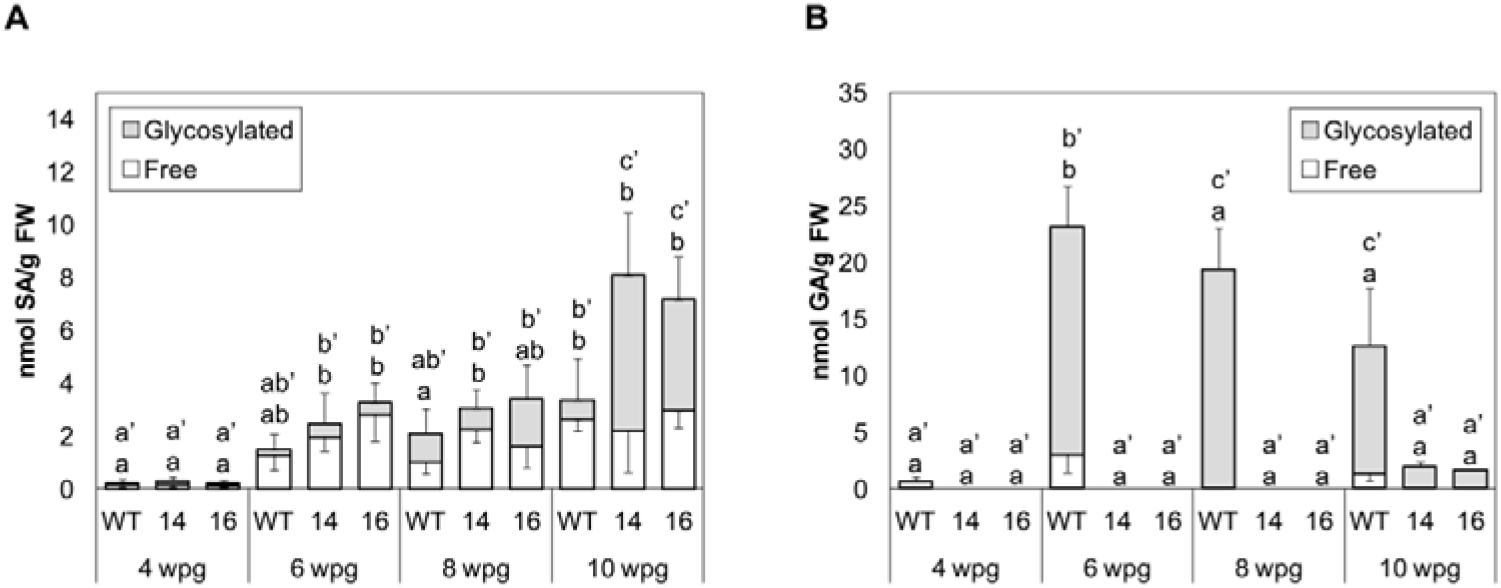
Evolution of free and glycosylated SA (A) and GA (B) accumulation in wild type (WT) and *RNAi_SlS5H* (lines 14 and 16) transgenic tomato plants. Samples were taken 4, 6, 8 and 10 weeks post-germination (wpg). Leaf extracts were analyzed by fluorescence HPLC. The data are presented as the mean ± standard deviation of a representative experiment (n=6). Significant differences for free and glycosylated SA and GA accumulation between different genotypes are represented by different letters since *p-*value < 0.05.

Taken together, our results appear to indicate that over-accumulation of SA in the *RNAi_SlS5H* transgenic tomato plants provokes a toxicity which may explain their lower growth rates and early senescence.

## DISCUSSION

Resistance to pathogens is associated with SA-mediated activation of plant immune response, being SA a key defence phytohormone with many potential derived applications in agriculture (Klessig et al., 2018). The fine-tuned regulation of SA accumulation constitutes an important point in plant immunity (Huang et al., 2018; Chen et al., 2019). This regulation can occur at different levels, including the control of its biosynthesis as well as its catabolism. Hydroxylation of SA to form 2,3-DHBA and GA can be considered part of its catabolism, but these forms also constitute a temporary source for SA, since their production could be reversible. Here we study the effect of the impairment of SA hydroxylation in tomato plants subjected to CEVd and *Pst* infections, which produce high and low levels of GA, respectively (Bellés et al., 1999; López-Gresa et al., 2016; López-Gresa et al., 2017; López-Gresa et al., 2019). Collectively, our findings provide insight into the modulation of SA metabolism upon biotic stress and add knowledge to previous results on this topic (Huang et al., 2018; Zhang and Li, 2019).

The SA-5 hydroxylation is performed by *AtS5H/DMR6* in *Arabidopsis thaliana*, displaying *s5h* mutants over accumulation of SA and enhanced resistance to *Pst* (Zhang et al., 2017). We have identified the tomato *S5H* ortholog and we studied its induction upon different pathogen infections which provoke SA and GA accumulation, including CEVd, TSWV, ToMV and a virulent and an avirulent strain of *Pst* (Figure 1). Similar results have been reported by Thomazella et al. (2021) from publicly available transcriptome data, including the bacteria *Xanthomonas gardneri* and *Pst*, the oomycete *Phytophthora capsici*, and the fungus *Moniliophthora perniciosa* as pathogens that trigger *SlS5H-DMR6-1* induction. Interestingly, our data suggest a correlation between *SlS5H* induction and symptomatology. Moreover, its induction correlates not only with symptom development but also with SA and GA accumulation in tomato, since both phenolics have been described to accumulate at very high levels in tomato plants infected by CEVd (Bellés et al., 1999; López-Gresa et al., 2016; López-Gresa et al., 2017; López-Gresa et al., 2019) which precisely provokes a high induction of *SlS5H*. According to that, *SlS5H* was also found to be induced by its own substrate at 6 hpt (Figure S2A), thus indicating a SA-mediated regulation of *SlS5H* expression. Whether that regulation is direct or may occur through the participation of different molecular elements remains unclear. This indirect regulation occurs for the transcription factor from apple *MdWRKY17*, which bounds to *MdDMR6* promoter and induces its expression (Shan et al., 2021). Our results reinforce the role of *SlS5H*, also known as *SlDMR6-1*, in SA metabolism.

SlDMR6-1 was described to be a 2-oxoglutarate-Fe(II) oxygenase acting on SA to produce GA *in vitro* (Thomazella et al., 2021). Here we confirm its *in vivo* SA 5-hydroxylase activity, by transiently overexpressing *SlS5H* in *Nicotiana benthamiana*. Plants agro-inoculated with a *35S:SlS5H* construction and further embedded with SA displayed lower levels of free and total SA, confirming that this phytohormone is a substrate for SlS5H *in vivo*, although a GA over-production was not detected (Figure S3). These results agree with those previously described in SA-treated *Nicotiana tabacum* plants which did not display GA accumulation (Bellés et al., 1999), probably indicating than this GA is rapidly catabolized in SA-embedded plants belonging to the *Nicotiana* genus. Our results confirm the extraordinary versatility of SA metabolism in different species, displaying levels of SA and GA accumulation up to 100-fold (Zhang et al., 2017).

Given the quantitative importance of GA accumulation in CEVd-infected tomato plants (Bellés et al., 1999; López-Gresa et al., 2016; López-Gresa et al., 2017; López-Gresa et al., 2019), we decided to study the role of *SlS5H* in this interaction by analysing the phenotype of *RNAi_SlS5H* transgenic tomato plants upon viroid infection. *SlS5H* silencing produced an enhanced resistance to CEVd, reducing both symptoms and viroid accumulation (Figure 2). Our results further extend the described broad spectrum resistance mediated by inhibition of SA hydroxylation (Thomazella et al., 2021) to a non-coding, RNA pathogen.

This resistance observed in *SlS5H*-silenced plants was accompanied by an induction of the SA-mediated plant defence response, since *PR1* expression leaves were higher in viroid infected *RNAi_SlS5H* transgenic tomato plants when compared to the corresponding infected non-transgenic plants (Figure 2E). In contrast, activation of the silencing mechanisms through *DCL1* and *DCL2* was reduced in *RNAi_SlS5H* transgenic tomato plants (Figure S4A and S4B), thus indicating these mechanisms are related to the viroid progression, which is limited in the transgenic plants, but not to the SA accumulation itself. In contrast, a correlation between SA accumulation and activation of silencing mechanisms has been previously described (Campos et al., 2014a), thus suggesting a role of SA in plant silencing. Besides, JA response appeared to be down-regulated in the *RNAi_SlS5H* transgenic tomato plants, displaying a lower induction of *TCI21* upon CEVd infection and wounding (Figures S4D and 8A), and showing higher susceptibility to *Botrytis cinerea* (Figure 9). The reciprocal antagonism between JA and SA (Thaler et al., 2012) and the higher induction of *PR1* in the transgenic lines points to a SA overaccumulation, which could be responsible for the observed SA-mediated plant resistance to CEVd. According to this, *RNAi_SlS5H* transgenic tomato plants displayed higher levels of total SA and significantly lower levels of total and free GA, confirming SlS5H activity *in vivo*. The enhanced resistance of *Sldmr6-1* lines to different classes of pathogens, such as bacteria, oomycetes and fungi has also been correlated with increased SA levels and transcriptional activation of immune responses (Thomazella et al., 2021).

Tomato plants infected with *Pst* have been described to display a different pattern of accumulation of both SA and GA when compared to CEVd infected tomato plants, reaching SA levels less than 10 nmol/g, and around 40 nmol/g for GA (López-Gresa et al., 2017), much lower than those detected upon CEVd infection (Bellés et al., 1999; López-Gresa et al., 2016; López-Gresa et al., 2017; López-Gresa et al., 2019). To test the effect of *SlS5H* impairment on the tomato response to bacteria, similar studies to those performed for CEVd were carried out. Our results were in accordance with the enhanced *PR1* induction (Figure 4C), enhanced resistance (Figure 4A), and reduced GA accumulation in *RNAi_SlS5H* transgenic tomato plants triggered by bacteria (Figure 5B). Unlike CEVd infection, an increase in SA accumulation was not detected in transgenic tomato plants upon *Pst* infection, thus indicating that SA catabolism is dependent on the tomato-pathogen interaction (Figure 5A).

The availability of different SA-deficient mutants and *NahG* transgenic plants in *Arabidopsis thaliana* background has provided vast information on the SA metabolism in response to bacterial infection in this species (Ding and Ding, 2020; Zeier, 2021). However, fewer studies have been done on the role of SA in tomato plants in response to bacteria. Therefore, to better understand the modulation of SA metabolism by *Pst* infection in tomato, and to compare it with that activated by CEVd, metabolomic analyses were performed. Our metabolomic assay confirmed that the glycosylated form of SA was the most accumulated compound in the *RNAi_SlS5H* transgenic tomato upon viroid infection (Figure 6B), whereas GA glycosylated form is described to be the most accumulated compound in CEVd-infected WT plants (López-Gresa et al., 2019), thus confirming the proposed SA hydroxylation *in vivo* activity for SlS5H. Strikingly, the impairment of SA hydroxylation in tomato plants uncovered alternative SA homeostasis upon *Pst* infection, since *RNAi_SlS5H* transgenic plants over-accumulated feruloyldopamine, feruloylquinic acid, feruloylgalactarate and 2-hydroxyglutarate (Figure 6C).

Feruloyldopamine is a hydroxycinnamic acid amide (HCAA) known to accumulate in tomato plants infected with a high *Pst* bacterial titre, thus producing a hypersensitive-like response (Zacarés et al., 2007; López-Gresa et al., 2011). It is worthy to note that this HCAA accumulation was accompanied by a rapid and sharp production of SA. These results appear to indicate that SA over-accumulation produced by bacterial infection could trigger the accumulation of this HCAA to prevent a toxic SA effect. The acyl-quinic acids are a diverse group of plant-derived compounds produced principally through esterification of an hydroxycinnamic acid and quinic acid (Clifford et al., 2017). Particularly, the flavonoid feruloylquinic acid has been described to accumulate in barley leaves upon ultraviolet and photosynthetically active radiation suggesting a protective role (Klem et al., 2015). In *Coffea canephora*, feruloylquinic acid is accumulated in juvenile leaves associated with chloroplasts, therefore suggesting a protective role against photooxidative damage (Mondolot et al., 2006). Feruloylgalactarate results from the reaction of feruloyl-CoA with galactaric acid. The accumulation of hydroxycinnamic acids esters of quinic acid and glucaric acids has also been described in tomato plants infected with *Ralstonia solanacearum*, the causal agent of bacterial wilt, a highly destructive bacterial disease (Zeiss et al., 2019). Finally, the accumulation of 2-hydroxyglutarate has been proposed to be linked to light-dependent photorespiration, it is related to oxidative stress and is considered as a marker for senescence in plants (Kuhn et al., 2013; Sipari et al., 2020). In this respect, the expression of 2-hydroxyglutarate dehydrogenase increases gradually during developmental and dark-induced senescence in *Arabidopsis thaliana* (Engqvist et al., 2011).

The accumulation of all these compounds may be somehow related with photooxidative damages, caused by biotic and abiotic stresses or senescence. There is a well-known interplay between SA and reactive oxygen species (ROS), being ROS signals involved both upstream and downstream SA signalling (Herrera-Vásquez et al., 2015). Therefore, the observed over-accumulation of these metabolites in the *RNAi_SlS5H* plants upon bacterial infection could be exerting a protective role against the photooxidative damage provoked by the transient over-accumulation of SA which could be promoting ROS production, as previously described (Herrera-Vásquez et al., 2015). On the other hand, there is a biosynthetic connection between SA and these phenolic compounds, displaying age-related differences in their biosynthesis in *Solanum lycopersicum* cv. *amateur* infected by *Pst* (Dadáková et al., 2020). Therefore, the accumulation of feruloyldopamine, feruloylquinic acid and feruloylgalactarate could also be the result of the SA catabolism.

The specific pathogen-triggered SA metabolism observed in this study, was accompanied with noticeable differences in the pattern of induction of SA biosynthesis genes provoked by the CEVd and *Pst* (Figure 10). Regarding *ICS* and *EPS1* expression levels, a significant induction was observed in WT plants after viroid and bacterial infection, respectively. However, no differences were observed after CEVd and *Pst* infection in transgenic plants when compared with mock controls. *ICS* works as an isochorismate synthase in the plastids, being required for SA biosynthesis (Ding and Ding, 2020). The impairment of its induction in transgenic plants upon CEVd infection appears to indicate a negative feed-back produced by SA over-accumulation (Figure 7A). However, this negative feed-back was not detected neither in transgenic plants upon bacterial infection, where the levels of SA were lower than after a viroid infection (Figure 8A), nor at *EPS1* level, which is described to act at the cytoplasm (Zeier, 2021). *EPS1* was significantly induced by *Pst* in WT tomato plants, while *RNAi_SlS5H* plants did not display that induction. In the same way, a reduction of the *PAL* expression was observed in transgenic plants upon CEVd infection coinciding with SA over-accumulation, and no differences in *PAL* induction were detected in tomato-*Pst* interaction. Regarding WT tomato plants, *PAL* was not induced by any pathogen, indicating that the main source of SA upon infection is also dependent on the IC pathway in tomato plants, as previously described in Arabidopsis (Wildermuth et al., 2001; Zhang and Li, 2019).

Finally, the *SAMT* regulation was opposite depending on the attack of the different studied pathogens in WT plants. While viroid caused a significant *SAMT* repression, bacterial infection produced *SAMT* induction. The reduced expression of *SAMT* upon CEVd inoculation could be due to the lower amount of its substrate which is redirected to form SAG (Figures 3A and 6B), since a correlation between SA and MeSA accumulations has been previously described (Hernández-Aparicio et al., 2021). However, the *SAMT* induction in tomato plants upon bacterial infection matches with the increase in the production of methyl salicylate (MeSA), salicylaldehyde, and ethyl salicylate described in this virulent plant-pathogen interaction (López-Gresa, 2017). The lower induction of *SAMT* upon *Pst* infection in transgenic plants could also be due to the lower amount of its substrate. Together, these results confirm the versatility of the SA metabolism in response to different pathogens.

These specific differences in tomato SA metabolism generated by both pathogens could also be explained by SA bacterial metabolism itself. Whilst CEVd is a non-coding pathogen totally dependent on host transcriptional machinery, *Pst* possesses its own SA biosynthetic pathway which is different from plants (Mishra and Baek, 2021). Moreover, plant SA accumulation is a target of suppression by *Pst* (Wilson et al., 2014), and the bacterial wilt pathogen *Ralstonia solanacearum* has been described to degrade plant SA in order to protect itself from inhibitory levels of this compound as well as to enhance its virulence on plant hosts (Lowe-Power et al., 2016). Therefore, while the observed SA metabolism caused by CEVd infection is totally plant-dependent, differences observed in *Pst*-infected *RNAi_SlS5H* tomato plants, including the lack of SA accumulation detected, could also be the result of the participation of the *Pst* SA metabolism.

The role of SA on leaf senescence has long been described (Morris et al., 2000). In this sense, impairment of S5H in Arabidopsis provokes an advanced senescent response (Zhang et al., 2013; Zhang et al., 2017). However, no clear phenotypic differences regarding senescence were described in *SlDMR6-1* tomato mutant (Thomazella et al., 2021). Here we have studied the effect of *SlS5H* silencing on the development of tomato plants, observing an early senescence phenotype provoked by SA over-accumulation (Figure 10). These differences could be due to the tomato variety employed for the studies, since Fla. 8000 was used by Thomazella et al (2021) and Moneymaker tomato plants were used in our studies. Alternatively, the differences between CRISPR knock out and RNAi silencing approaches could also be considered. According to our results, a higher induction of both *SAG12* and *NOR* senescence markers (Ma et al., 2019), as well as *PR1*, a molecular marker for the defence response (Conejero, V., Bellés, J. M., García-Breijo, F., Garro, R., Hernández-Yago, J., Rodrigo, I., & Vera, 1990), was detected in both transgenic plants silencing *SlS5H* (Figure 10). As previously stated, the toxic effect of SA over-accumulation, also responsible for the observed cell death and the decrease in chlorophyll, could be related with ROS accumulation (Herrera-Vásquez et al., 2015). Our results provide genetic evidence that *SlS5H* has an important role in regulating the onset and rate of leaf senescence in tomato by fine-tuning the endogenous levels of SA.

Because of the balance between *de novo* biosynthesis, catabolism and reversible deactivation of SA, plants manage to maintain their endogenous levels of SA. Here we demonstrate that there is an additional complexity level caused by specific pathogen-induced SA homeostasis, suggesting that the framework of SA biology established in *Pst*-infected *Arabidopsis thaliana* plants should be reconsidered for each specific plant-pathogen interaction.

## MATERIALS AND METHODS

### Plant materials and growth conditions

Tomato Rio Grande plants, containing the *Pto* resistance gene, were used to establish the virulent and avirulent interaction upon bacterial infections. In the rest of experiments, transgenic tomato (*Solanum lycopersicum*) plants silencing the endogenous salicylate 5-hydroxylase gene (*SlS5H*) and the cultivar Moneymaker, the isogenic parental line of *RNAi_SlS5H*, were used.

Tomato seeds were surface sterilized with sodium hypochlorite. After sterilization, seeds were sown in 12 cm-diameter pots and grown in standard greenhouse conditions, with a temperature between 25-30 °C, a relative humidity of 50-70 % and long day photoperiod (16 h light/8 h darkness).

For transient expression experiments, *Nicotiana benthamiana* plants were cultivated in the same conditions as tomato plants.

### Vector construction

The full-length cDNA (1014 bp) of salicylate 5-hydroxylase gene (*SlS5H;* Solyc03g080190) was amplified by RT-PCR from leaves of Moneymaker tomato plants infected with CEVd, 4 weeks after inoculation, using 5′ ATGGAAACCAAAGTTATTTC-3′ as the forward primer and 5′-GTTCTTGAAAAGTTCCAAAC-3′ as the reverse primer. The resulting PCR product was cloned into the pCR8/GW/TOPO entry vector (Invitrogen), following the manufacturer’s protocol, and was sequenced. Then *SlS5H* was subcloned in the pGWB8 Gateway binary vector (Nakagawa et al., 2007). This vector carries the CaMV35S promoter and the hexahistidine tag (6XHis) which is attached to the C-terminus of the recombinant protein.

In order to generate the *SlS5H*-silenced transgenic tomato plants, the method described by Helliwell and Waterhouse was followed (Helliwell and Waterhouse, 2003). Briefly, a selected 400 bp sequence of *SlS5H* was amplified from the full-length cDNA clone using the forward primer 5′-GGCTCGAGTCTAGAGGGAAATTCGTCAA-3′, which introduced restriction sites *Xho*I and *Xba*I, and the reverse primer 5′-CCGAATTCGGATCCACCGTTACTTTACTGC-3′, which added restriction sites *BamH*I and *EcoR*I. The PCR product was first cloned in the pGEM T Easy vector (Promega) and sequenced. After digestion with the appropriate restriction enzymes and purification, the two *SlS5H* fragments were subcloned into the pHANNIBAL vector in both the sense and the antisense orientations. Finally, the constructs made in pHANNIBAL were subcloned as a *Not*I flanked fragment into pART27 binary vector to produce highly effective intron-containing “hairpin” RNA silencing constructs (Gleave, 1992). This vector carries the neomycin phosphotransferase gene (NPT II) as a transgenic selectable marker.

### *N. benthamiana* agroinfiltration and tomato transformation

The pGWB8-SlS5H construction and the pGWB8 empty vector were transformed into the *Agrobacterium tumefaciens* C58 strain, while the pART27-SlS5H construction was transformed into *A. tumefaciens* LBA4404. Leaves of 4-week-old *N. benthamiana* plants were infiltrated with the *A. tumefaciens* C58 strain carrying pGWB8-SlS5H or the empty vector, along with a 1:1 ratio of the C58 strain carrying the p19 plasmid, which encodes the silencing suppressor protein p19 (Lakatos et al., 2004). Tomato Moneymaker cotyledons were co-cultured with *A. tumefaciens* LBA4404 carrying the pART27-SlS5H construction to generate the RNAi *SlS5H*-silenced transgenic tomato plants (*RNAi_SlS5H*). The explant preparation, selection and regeneration methods followed those published by Ellul and co-workers (Ellul et al., 2003). The tomato transformants were selected in kanamycin-containing medium and propagated in soil. Moneymaker wild-type tomato plants regenerated *in vitro* from cotyledons under the same conditions as the transgenic lines were used as controls in subsequent analyses. The transgenic plants generated in this study have been produced, identified and characterized in our laboratory and are to be used exclusively for research purposes.

### Production of S5H recombinant protein in *Nicotiana benthamiana*

Agroinfiltration of *N. benthamiana* leaves with *Agrobacterium tumefaciens* C58, carrying either *pGWB8-SlS5H* or *pGWB8* empty vector was performed according to Yang et al. (Yang et al., 2000). Three days after the agro-inoculation, plants were embedded in a solution of 1 mM SA (see SA treatments) and samples were collected 24 upon the treatment. For western blot analysis, 5 g of frozen agroinfiltrated *N. benthamiana* leaves were ground and resuspended in 1 mL of extraction buffer (50 mM Tris-HCl, pH 7.5, containing 15 mM 2-mercaptoethanol). Proteins were separated by SDS-PAGE and stained with Coomassie Brilliant Blue R-250 (Conejero and Semancik, 1977) or transferred to nitrocellulose filters (OPTITRAN, Schleicher&Schuell) following the protocol described by Towbin et al. (Towbin et al., 1979). S5H recombinant protein was examined by using specific anti-His mouse antibody (Novagen) as described in (López-Gresa et al., 2016).

### SA treatments

The SA treatments were carried out by stem-feeding (Campos et al., 2019). Four-week-old tomato plants or agro-inoculated *N. benthamiana* plants were excised above cotyledons, and stem cuts were immediately immersed in a 2 mM or 1 mM SA solution, respectively. After 30 min, all the stems were transferred to water, and leaf samples from three biological replicates were taken at 0, 0.5, 1, 6, 24 and 48 h post-treatment.

### Viroid Inoculation

For these assays, tomato plants were grown on a growth chamber with a temperature between 28 °C/24 °C and a relative humidity of 60%/85% (day/night). Viroidal inoculum was prepared from leaves of CEVd-infected tomato plants as previously described (Semancik et al., 1988) and the first cotyledon and leaf of each plant were inoculated using carborundum as abrasive. Mock plants were inoculated with sterile water. Disease and symptom severity was recorded periodically. Leaf samples from six infected or mock-inoculated plants were harvested at 2- and 3-weeks post inoculation (wpi).

### *Pseudomonas syringae* inoculation

The bacterial strain used in this study was *Pseudomonas syringae* pv. *tomato* DC3000 (*Pst*). For incompatible interaction, the infection was performed by the bacterial strain Pst DC3000 that contains deletions in genes *avrPto* and *avrPtoB* (Pst DC3000 ΔavrPto/ΔavrPtoB) (Ntoukakis et al., 2009)

Pathogen inoculation was performed in 4-week-old tomato plants by immersion, as previously described (López-Gresa et al., 2018). For mock treatments, plants were immersed in 10 mM sterile MgCl_2_ containing 0.05% Silwet L-77. The third and fourth leaves from six plants per treatment and genotype were harvested 24 h after inoculation.

### *Botrytis cinerea* inoculation assays

The *B. cinerea* strain used was CECT2100 (Spanish Type Culture Collection). Fungal hyphae were grown on potato dextrose agar for 14 days at 24 °C in darkness. Spore suspensions were prepared by scraping surface plates, washing with sterile water, and filtering through cotton. Finally, the concentration was adjusted to 10^6^ spores/mL. Three leaflets per plant were spotted with a 5 mL droplet of the spore suspension. All the experiments were carried out into inoculation chambers to maintain the proper high humidity conditions. Photographs, lesion size measurements and sampling were performed at 5 days after inoculation.

### Chlorophyll Content Determination

Chlorophyll quantification was carried out by the method of Arnon (Arnon, 1949). Frozen leaf tissue was homogenized with 80% acetone and then incubated overnight at 4 °C. Then, samples were centrifuged and the absorbance of the extracted solution was measured at 645 and 663 nm. Determination of chlorophyll a, b and total concentration was determined according to Arnon’s equations.

### Extraction and HPLC analysis of salicylic and gentisic acids

Extraction of free and total SA and GA from tomato leaflets was performed according to our previously published protocol (Bellés et al., 2006). Aliquots of 30 μL were injected through a Waters 717 autosampler into a reverse-phase Sun Fire 5-mm C18 column (4.6 mm x 150 mm) equilibrated in 1% (v/v) acetic acid at room temperature. A 20-min linear gradient of 1% acetic acid to 100% methanol was applied using a 1525 Waters Binary HPLC pump at a flow rate of 1 mL/min. SA and GA were detected with a 2475 Waters Multi-l Fluorescence detector (λ excitation 313 nm; λ emission 405 nm) and were quantified with the Waters Empower Pro software using authentic standard compounds (SA sodium salt and GA, Sigma–Aldrich, Madrid, Spain). Standard curves were performed for each compound using similar concentration ranges to those detected in the samples. Data were corrected for losses in the extraction procedure, and recovery of metabolites ranged between 50 and 80%.

### UPLC-ESI-QTOF-MS analysis

For UPLC-MS analysis, frozen tomato leaves (100 mg) were ground into powder in liquid nitrogen and extracted in 1 mL of methanol/water (80:20, v/v) for chromatographic analysis.

UPLC separations were performed on a reverse phase Poroshell 120 EC-C18 column (3 × 100 mm, 2.7 μm) (Agilent Technologies) operating at 30 °C and a flow rate of 0.4 mL/min. The mobile phases used were acidified water (0.1 % formic acid) (Phase A) and acidified acetonitrile (0.1 % formic acid) (Phase B). Compounds were separated using the following gradient conditions: 0–10 min, 1–18 % phase-B; 10–16 min, 18–38 % phase-B; 16–22 min, 38– 95 % phase-B. Finally, the phase B content was returned to the initial conditions (1 % phase-B) for 1 min and the column re-equilibrated for five minutes more. 7 μL of the sample was injected using flow through needle (FTN) injection with a 15 mL syringe. The sample compartment in the auto sampler was maintained at 7.0 °C.

The UPLC system was coupled to a quadrupole-time-of-flight (maXis Impact HR Q-ToF-MS, (Bruker Daltonik GmbH, Bremen, Germany) orthogonal accelerated Q-ToF mass spectrometer, was performed using HR-ToF-MS in negative electrospray ionization mode using broadband collision induced dissociation (bbCID). High and low collision energy data were collected simultaneously by alternating the acquisition between MS and bbCID conditions.

Parameters for analysis were set using negative ion mode, with spectra acquired over a mass range from 50 to 1200 *m/z*. The optimum values of the ESI-MS parameters were: capillary voltage, -4.0 kV; drying gas temperature, 200 °C; drying gas flow, 9.0 L/min; nebulising gas pressure, 2 bars; collision RF, 150 Vpp; transfer time 72 μs, and pre-pulse storage, 5 μs.

At some stage in the UHPLC method development, an external apparatus calibration was performed using a KD Scientific syringe pump (Vernon Hills, IL) directly linked to the interface, passing a solution of sodium formate with a flow rate of 180 μL/h. The instrument was calibrated externally before each sequence with a 10 mM sodium formate solution.

Using this method, an exact calibration curve based on numerous cluster masses each differing by 68 Da (CHO2Na) was obtained. Due to the compensation of temperature drift in the Q-TOF, this external calibration provided accurate mass values for a complete run without the need for a dual sprayer set up for internal mass calibration.

### Non-targeted metabolomics analysis and quality control

All samples were injected in the same batch and the order of sample injection was randomized to avoid sample bias. A mixture with one replicate of each group of samples was used as ‘‘quality control’’ (QC) and was injected at the beginning, in the middle and at the end of the batch. Besides, methanol/water injections were included every five samples as a blank run to avoid the carry-over effect.

For the untargeted analysis of the polar and semi-polar profiles, the QToF-MS data were processed with XCMS online resources (https://xcmsonline.scripps.edu) with the appropriate script for the alignment of chromatograms and the quantification of each MS feature (Rambla et al., 2015). The resulting dataset was submitted to a Principal Component Analysis (PCA) study by the SIMCA-P software (v. 11.0, Umetrics, Umeå, Sweden) using unit variance (UV) scaling.

Metabolite identification was based on comparison of accurate mass, retention time, MS/MS fragments and CCS values with online reference databases including Respect (https://spectra.psc.riken.jp/), Metlin (https://metlin.scripps.edu/), HMDB (https://hmdb.ca/), Lipidmap (https://www.lipidmaps.org/), in-house databases based on commercial standards and theoretical MS/MS tags, and bibliographies. The CCS value acceptable error was <5% with MS tolerance of 5 p.p.m., and MS/MS tolerance of <10 mDa, at least one major fragment was found.

### RNA extraction and quantitative RT-PCR analysis

The total RNA of tomato leaves was extracted using the TRIzol reagent (Invitrogen, Carlsbad, CA, United States), following the manufacturer’s protocol. RNA was then precipitated by adding one volume of 6 M LiCl and keeping it on ice for 4 h. Afterward the pellet was washed using 3 M LiCl and was dissolved in RNase-free water. Finally, to remove any contaminating genomic DNA, 2 U of TURBO DNase (Ambion, Austin, TX, United States) were added per microliter of RNA. For the quantitative RT-PCR (qRT-PCR) analysis, one microgram of total RNA was employed to obtain the corresponding cDNA target sequences using an oligo(dT)18 primer and the PrimeScript RT reagent kit (Perfect Real Time, Takara Bio Inc., Otsu, Shiga, Japan), following the manufacturer’s directions. Quantitative PCR was carried out as previously described (Campos et al., 2014b). A housekeeping gene transcript, actin or elongation factor 2, was used as the endogenous reference. The PCR primers were designed using the online service Primer3 (https://primer3.ut.ee/) and are listed in Table S1.

**Table S1.**
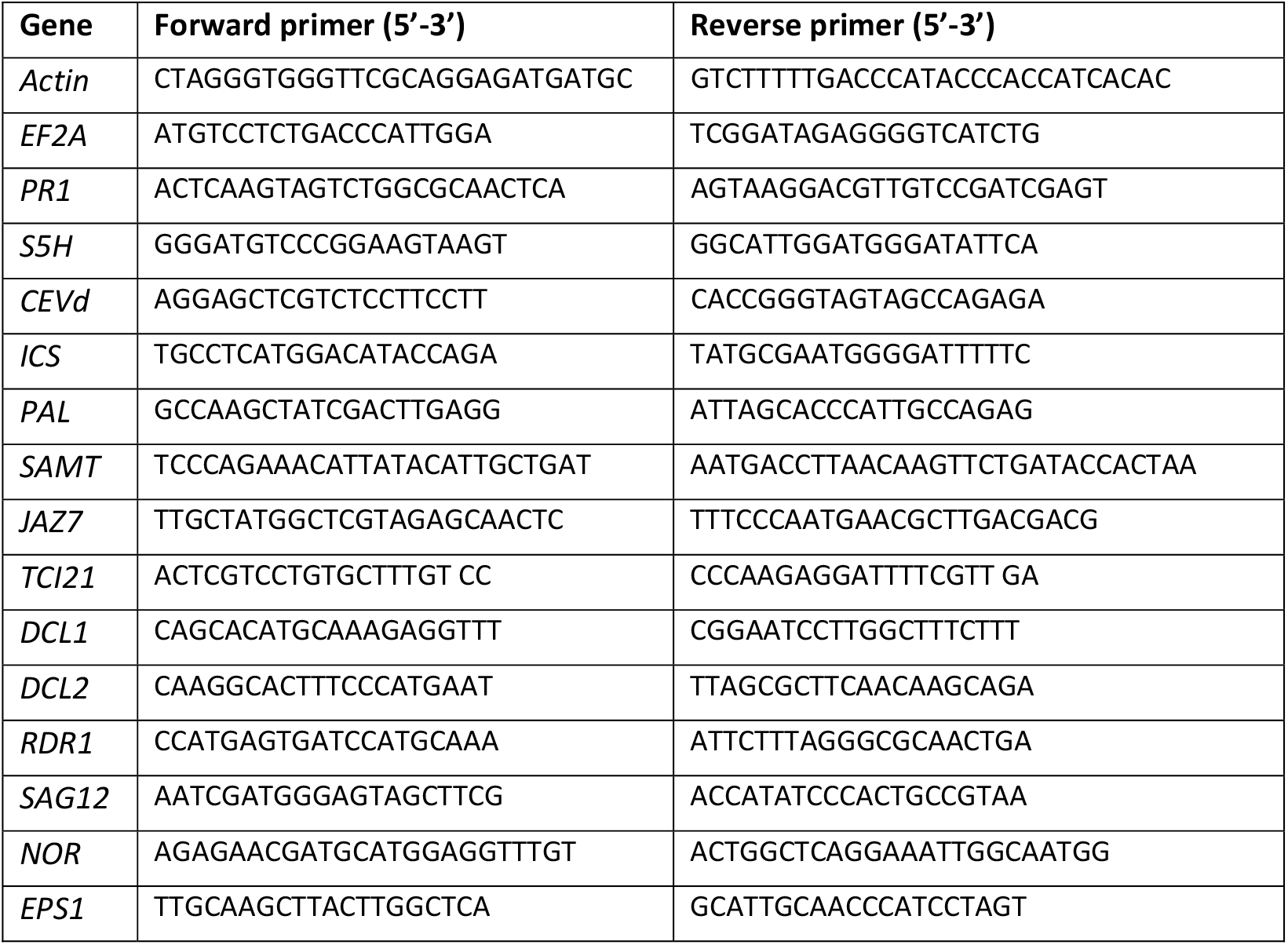
Primer sequences used for quantitative RT-PCR analysis of the selected tomato genes.

### Statistical analysis

The statistical analysis of two or more variables was carried out by using Student’s *t*-test or analysis of variance (ANOVA), respectively, employing the Prism 9 software (https://www.graphpad.com/). In all the analyses, a *p*-value < 0.05 was considered statistically significant.

## ACCESSION NUMBERS

Sequence data from this article are found in the GenBank/EMBL and Solgenomics data libraries under the following accession numbers: *Actin* (Solyc01g104770); *EF2A* (NM_001321725); Solyc03g080190 (*SlS5H*); *PR1* (X71592); *ICS* (Solyc06g071030); *JAZ7* (Solyc11g011030); *TCI21* (Solyc03g098790); *DCL1* (Solyc10g005130); *DCL2 (*Solyc11g008540); *RDR1* (Solyc05g007510); *SAG12* (AT5G45890 tomato ortholog; Solyc02g076910); *NOR* (NM_001247723.2); *EPS1* (Solyc08g005890); *PAL* (NM_001320040).

## ACKNOWLEDGMENTS

We would like to thank the “Metabolomics Platform of CEBAS-CSIC” (Centro de Edafología y Biología Aplicada del Segura, Murcia, Spain), especially to Dr. José E. Yuste (Technical Coordinator) for his excellent support in the phenolics identification of this study.

## FIGURE LEGENDS

**Figure S1. Phylogenetic analysis of *AtS5H* orthologs in tomato**. The box in the phylogenetic tree highlights *AtS5H* from *Arabidopsis thaliana* (At5g24530) and its closest homolog in tomato (Solyc03g080190). The multiple alignment was made using ClustalW and the dendrogram was built using the MegAlign program from the Lasergene package (DNASTAR, Madison, Wisconsin, USA).

**Figure S2. SA-induced expression of *SlS5H* in wild type (WT) and *RNAi_SlS5H* transgenic tomato lines 14 and 16**. *SlS5H* **(A)** and *PR1* **(B)** expression of tomato plants treated with 2 mM of SA (SA) or water (MOCK) by stem feeding at 0, 0.5, 1, 6, 24 and 48 hours post-treatment. The qRT-PCR values were normalized with the level of expression of the actin gene. The expression levels correspond to the mean ± the standard error of a representative experiment (n=3). Significant differences between mock and infected or treated plants at different time points are represented by different letters when *p-*value < 0.05.

**Figure S3. S5H *in vivo* activity in *Nicotiana benthamiana* plants. (A)** SDS-PAGE (left panel) and western blot analysis (right panel) of *N. benthamiana* plants agroinoculated with pGWB8 empty vector (C) or pGWB8-SlS5H (S5H). **(B)** Representative image of the cloning cassette. Nanomoles of SA **(C)** and GA **(D)** per gram of fresh weight in *Nicotiana benthamiana* leaves embedded with SA and agroinoculated with the construction pGWB8-S5H, compared with its control (plasmid pGWB8 without insert). The results correspond to a representative experiment (n=3). Student’s *t*-statistic analysis shows the mean ± standard deviation. No statistical differences were observed. 2,3-DHBA levels were not detected.

**Figure S4. Gene expression analysis of wild type (WT) and *RNAi_SlS5H* (lines 14 and 16) transgenic tomato plants, mock-inoculated (MOCK) and inoculated with CEVd (CEVd)**. *DCL1* **(A)**, *DCL2* **(B)**, *RDR1* **(C)** and *TCI21* **(D)** gene expression was analyzed 3 weeks after viroid infection. The qRT-PCR values were normalized with the level of expression of the actin gene. The expression levels correspond to the mean ± the standard error of a representative experiment (n=3). The significant differences between different genotypes and infected or mock-inoculated plants are represented by different letters since *p-*value < 0.05.

**Figure S5. Score plot of PCA based on whole range of on the whole array of the mass spectra within a *m/z* range from 100 to 1500 using unit variance (UV) scaling method of methanolic extracts from tomato leaves. (A)** CEVd infected plants at 3 wpi, green: wild type (WT); light purple: *RNAi_SlS5H* 14; dark purple: *RNAi_SlS5H* 16; **(B)** *Pst* infected plants at 24 hpi, yellow: wild type (WT); orange: *RNAi_SlS5H* 14; brown: *RNAi_SlS5H* 16.

**Figure S6. Analysis of phenotypic differences between WT and *RNAi*_S5H transgenic lines 14 and 16**. Differences related to weight **(A)**, conductivity **(B)** and chlorophyll content **(C)** in WT and *RNAi_SlS5H* 14 and 16 transgenic plants were measured 10 weeks after germination. Bars represent the mean ± the standard deviation of a representative experiment (n=6).

## Notes

**Funding information** This work was supported by Grant PID2020-116765RB-I00 funded by MCIN/AEI/ 10.13039/501100011033 and Grant AICO/2017/048 from the Generalitat Valenciana. Work in the lab is also supported by grant PROMETEU/2021/056 by Generalitat Valenciana. C.P. was a recipient of a predoctoral contract of the Generalitat Valenciana (ACIF/2019/187).

### Competing Interest Statement

The authors have declared no competing interest.

